# Tertiary lymphoid structures are associated with enhanced macrophage activation, immune checkpoint expression and predict outcome in cervical cancer

**DOI:** 10.1101/2023.08.17.552583

**Authors:** Laurent Gorvel, Marylou Panouillot, Marie-Sarah Rouvière, Emilien Billon, Stéphane Fattori, Jumaporn Sonongbua, Nicolas Boucherit, Amira Ben Amara, Olivia Quilichini, Samuel Granjeaud, Clara Degos, Jacques A. Nunes, Xavier Carcopino, Eric Lambaudie, Anne-Sophie Chretien, Renaud Sabatier, Marie-Caroline Dieu-Nosjean, Daniel Olive

## Abstract

**Background:** Cervical tumors are usually treated using surgery, chemotherapy, and radiotherapy, and would benefit from immunotherapies. However, the immune microenvironment in cervical cancer remains poorly described. Tertiary lymphoid structures (TLS) were recently described as markers for better immunotherapy response and overall better prognosis in cancer patients.

**Methods:** We integratedly evaluated the cervical tumor immune microenvironment, and specifically TLS importance, using combined high-throughput phenotyping, soluble factor dosage, spatial interaction analyses, and statistical analyses.

**Results:** We demonstrate that TLS presence is associated with a more inflammatory soluble microenvironment, with the presence of B cells as well as more activated macrophages and dendritic cells (DCs). Furthermore, this myeloid cell activation is associated with expression of immune checkpoints, such as PD-L1 and CD40, and close proximity of activated conventional DC2 to CD8^+^ T cells, therefore indicating better immune interactions and tumor control. Finally, we associate TLS presence, greater B cell density, and activated DC density to improved progression-free survival, and present it as an additional prognostic marker.

**Conclusion:** To conclude, our results provide an exhaustive depiction of the cervical tumor immune microenvironment where TLS presence marks cell activation and immunotherapy target expression. These findings provide predictive clues for patient response to targeted immunotherapies.

**Significance:** TLS maturation stratifies cervical cancer patients and associates with improved prognosis. TLS associate with the expression of immune checkpoints, notably in the macrophage compartment, which may represent a new therapeutic strategy.

## Introduction

Cervical cancer is the fourth most common cancer in women and is still a public health issue in developing countries (1). Cervical tumors are human papillomavirus (HPV)-positive in more than 95% of cases. Vaccination against HPV provides good protection against the development of cervical tumors; however, vaccination rates remain low worldwide (50% in both Europe and the USA)(1). Most cervical tumors comprise well-infiltrated squamous cell carcinomas (SCC), the rest being less-infiltrated adenocarcinomas (ADK)(2). Cervical SCCs are usually associated with better overall survival (OS) than ADKs (5-year OS rates of 80% and 60%, respectively). Tumor severity is defined by International Federation of Gynecology and Obstetrics (FIGO) stages, where a higher FIGO stage corresponds to worse severity, lymph node (LN) invasion and metastasis, and correlates with poorer OS (decreasing to 55% at 5 years for FIGO stage IVa)(3).

Cervical tumors are treated using surgery and brachytherapy and chemotherapy. These are debilitating treatments, especially for younger women in age to expect children. Immunotherapies provide an advantageous set of treatment as they often present less toxicities than chemotherapy or radiations. However, they require a well described immune environment, which is currently understudied in cervical tumors. It is known that the T cell compartment harbors regulatory T cells (Tregs), which are often associated with poor prognosis, and tumor-infiltrating conventional CD4^+^ and CD8^+^ T cells, including HPV-specific T cells (4-6). The myeloid compartment is less well studied, with research focusing on the Tumor-Associated Macrophage (TAM) infiltrate phenotype in the tumor bed and how this associate with survival. Overall, whether they express CD163, CD206, or CD204, TAMs are associated with advanced FIGO stages(7-10). Interestingly, the expression of programmed death-ligand 1 (PD-L1) by TAM is associated with programmed death-1 (PD-1) expression in the CD8^+^ and CD4^+^ T cell compartments and improves prognosis when the ratio is in favor of CD8^+^ T cell infiltrate (11-13). However, most of these studies only correlate presence or absence of specific cell types to prognosis or therapy response, and do not assess their spatial distribution within the tumor microenvironment.

Determining the localization and interactions of subsets of immune cells in the stroma, or within the tumor nests is now critical to understanding tumor-specific treatment responses. Indeed, the organization of the immune infiltrate, particularly into tertiary lymphoid structures (TLS), has been associated with improved prognosis and response to immunotherapy in cancer patients (14). TLS are organized aggregates of immune cells which develop at the site of chronic inflammation and display a similar organization to lymphnodes (15). Indeed, TLS exhibit a B cell zone that includes mainly germinal center B cells and follicular dendritic cells (FDCs), adjacent to a T cell zone comprising mainly T cells and activated DCs (aDCs)(16,17). As TLS are heterogeneous, they warrant further investigation in specific tumor microenvironments, as both their composition and the associated overall immune infiltrate may provide insights for tumor-specific therapies and biomarkers(14). The presence of TLS within cervical tumors has not been directly demonstrated yet and remains at the gene signature detection level(18). However, TLS presence in HPV-positive head and neck SCCs (HNSCCs) is associated with a higher B cell infiltrate and follicular helper T cells (Tfh)(19). A greater proportion of myeloid cells and CD8^+^ T cells in the vicinity of the TLS has also been reported. Together, these parameters are associated with an improved prognosis in patients with HPV-positive HNSCC. Among the gynecological cancers, endometrial tumors are reported to harbor TLS(20), and these are associated with improved prognosis in microsatellite instability (MSI) endometrial tumors, allowing stratification for immunotherapy within this patient group(20). Indeed, the improved prognosis seen in MSI patients suggests that the presence of TLS and its associated immune infiltrate should be added to the molecular classification of endometrial cancer(20).

Therefore, integrated investigation of the cervical tumor immune infiltrate is clearly warranted to understand the different cellular compartments and their interactions. Here, we present an in-depth study of TLS presence in the cervical cancer immune microenvironment and its impact on the overall immune infiltrate, at both the cellular and spatial levels, and how these parameters may contribute to overall prognosis and therapy response.

## Methods

### Human tumor samples

Human cervical tumor samples (n=34) were obtained from patients, who were included in the prospective XAC-03 (NCT02875990) and GC-Bio-IPC (NCT01977274) clinical trials, conducted from 2013 to 2023. Patient follow-up was performed until December 2023 by assessing the occurrence of relapses or death. The XAC-30 and GC-Bio_IPC trials were approved by the institutional review board (Comité d’Orientation Stratégique (COS), Marseille, France) of the Paoli-Calmettes Institute and the Assistance Publique des Hopitaux de Marseille (AP-HM), respectively. In accordance with the Declaration of Helsinki, written informed consent was obtained from all patients.

Tumor samples were acquired prior to patients receiving any treatment. Healthy cervixes were obtained from hysterectomies performed in cases of benign uterine pathologies (endometriosis and fibroids). Before sampling and fixation, the cervix was assessed by a pathologist to confirm it was normal. Clinical characteristics of the patients from whom samples were obtained are shown in **Table 1**. A list of patient samples used in each experiment is available in **Suppl. Table 9**.

**Table 1:** Clinical characteristics of cervical cancer patients.

|  |  | Tumor samples* | Healthy samples |
| --- | --- | --- | --- |
| Characteristic | - | n=34 | n=5 |
| Age, median (range) | Current (years) | 58 (37-80) |  |
|  | At diagnosis (years) | 53 (32-77) |  |
| Histology, n | SCC** | 28 | NA |
|  | ADK** | 5 | NA |
|  | Other** | 1 | NA |
| FIGO stages, n | IB1 | 4 | NA |
|  | IIA1 | 2 | NA |
|  | IIA2 | 1 | NA |
|  | IIB | 14 | NA |
|  | IIIA | 1 | NA |
|  | IIIB | 3 | NA |
|  | IIIC1 | 5 | NA |
|  | IVA | 2 | NA |
|  | IVB | 2 | NA |
| Emboli, n | Positive | 3 | 0 |
|  | Negative | 31 | 5 |
| LN invasion, n | Positive | 14 | 0 |
|  | Negative | 20 | 5 |
| Adverse events, n | Metastasis | 4 | 0 |
|  | Relapses | 5 | 0 |
|  | Deaths | 2 | 0 |
| Treatments, n | RCT*** + Barytherapy | 26 | NA |
|  | Radio-Chemotherapy | 5 | NA |
|  | Chemotherapy | 1 | NA |
|  | Barytherapy | 2 | NA |

\*Note: cervical tumors are considered 100% HPV+
\*\* Note: SCC: squamous cell carcinoma; ADK: adenocarcinoma; Other: mixed origins, serous, clear cell carcinomas
\*\*\*Note: RCT: Radio-chemotherapy

### Immunohistochemistry staining of FFPE slides

All reagents and antibodies used for multiplex immunohistochemistry assay are listed in **Suppl. Table 1–3**. After deparaffinization of two slides for e ach patient, epitope retrieval was performed with the Dako Target retrieval solution (Dako) at pH 6.0 in a water bath for 30 minutes (min) at 97°C. After washing with TBS (Euromedex), tissue sections were treated with 3% H_2_0_2_ at room temperature (RT) for 15 min to inactivate endogenous peroxidase. After washing, sections were incubated with Protein Block solution for 30 min at RT. To visualize TLS, tissue sections were incubated with anti-DC-Lamp and anti-CD3 antibodies for one slide, and with anti-CD20 and anti-CD21 antibodies for a second slide. After washing with TBS-0.04% Tween 20, the first slide was incubated with donkey F(ab)’_2_ anti-rat IgG-biotin and donkey F(ab)’_2_ anti-rabbit IgG-alkaline phosphatase (AP), and the second slide with goat IgG anti-mouse IgG2a-biotin and goat IgG anti-mouse IgG1-AP, for 30 min at RT. After washing steps, biotin-conjugated antibodies were revealed with a streptavidin-horseradish peroxidase (HRP) solution, incubated for 30 min at RT. CD3 and CD21 immunostaining revelations were performed by adding SAP solution; DC-LAMP and CD20 were revealed with AEC solution. Slides were coverslipped with Glycergel and scanned using a Nanozoomer scanner (Hamamatsu).

### Determination of TLS positive samples

As described above, FFPE slides were stained for CD3, CD20, CD21 and DC-LAMP. B cells (CD20+) inside B cell aggregates and activated DCs (DC-LAMP+) located in T cell (CD3+) aggregates were counted. Their density was assessed by dividing B cell or activated DC count by the surface of the Tumor (in µm²). The obtained B cell and activated DC densities (in count/µm²) were then scored by quartile, and combined to assess TLS presence. A higher score for B cell and activated DC density resulted in higher TLS score (quartile) and more formed TLS while a lower TLS score resulted in absence of TLS or unorganized aggregates. TLS presence and maturity were then confirmed by microscopy by a trained pathologist and CD21 staining marking Follicular DCs (FDCs) (**Suppl. Fig. 1, Table 2**).

**Table 2:** TLS scoring parameters and values.

| Patient | DC density<br>(cell/mm<sup>2</sup>) | B cell<br>density<br>(cell/mm<sup>2</sup>) | TLS counts | fDCs in<br>B cell zone | TLS<br>maturity<br>scoring |
| --- | --- | --- | --- | --- | --- |
| GC1282 | 8.05122E-05 | 0.000332744 | 33 | NO | IMMATURE |
| GC1322 | 1.8465E-05 | 0.000177513 | 15 | NO | IMMATURE |
| GC1471 | 0.00010261 | 6.02821E-05 | 13 | NO | IMMATURE |
| XACI11 | 2.87553E-05 | 0.000133947 | 17 | NO | IMMATURE |
| XACI16 | 2.75875E-05 | 0.000238909 | 16 | NO | IMMATURE |
| XACN12 | 4.92789E-05 | 0.000551579 | 22 | NO | IMMATURE |
| XACN14 | 8.10343E-05 | 0.000322344 | 9 | NO | IMMATURE |
| XACN2 | 3.68913E-05 | 4.13829E-05 | 31 | NO | IMMATURE |
| XACN3 | 0.000116616 | 7.35915E-05 | 25 | NO | IMMATURE |
| XACN7 | 4.69993E-05 | 0.000114184 | 6 | NO | IMMATURE |
| GC1389 | 8.07186E-05 | 0.000274351 | 48 | YES | MATURE |
| GC1468 | 1.26599E-05 | 0.000127847 | 59 | YES | MATURE |
| XACC1 | 3.64324E-05 | 0.000276118 | 64 | YES | MATURE |
| XACN11 | 2.53324E-05 | 0.000443208 | 29 | YES | MATURE |
| XACN15 | 6.09574E-05 | 6.25658E-05 | 19 | YES | MATURE |
| XACN8 | 9.5982E-05 | 0.001264506 | 12 | YES | MATURE |
| GC1323 | 2.89623E-06 | 2.13661E-05 | 3 | NO | NEGATIVE |
| GC1358 | 1.82409E-05 | 2.31198E-05 | 2 | NO | NEGATIVE |
| GC1364 | 3.59181E-07 | 0 | 0 | NO | NEGATIVE |
| GC1374 | 7.8166E-06 | 1.87018E-05 | 6 | YES | NEGATIVE |
| GC1393/XACI15 | 1.05998E-05 | 2.03246E-06 | 1 | NO | NEGATIVE |
| GC1409 | 1.95797E-05 | 0.000100016 | 1 | NO | NEGATIVE |
| GC1483 | 1.06908E-05 | 8.33479E-06 | 2 | NO | NEGATIVE |
| XACI06 | 0 | 0 | 0 | NO | NEGATIVE |
| XACI07 | 3.48074E-06 | 0 | 0 | NO | NEGATIVE |
| XACI14 | 0 | 0 | 0 | NO | NEGATIVE |
| XACN10 | 1.88341E-05 | 0 | 0 | NO | NEGATIVE |
| XACN13 | 2.3648E-06 | 2.38441E-06 | 1 | NO | NEGATIVE |
| XACN16 | 7.10688E-05 | 6.31852E-06 | 2 | NO | NEGATIVE |
| XACN4 | 6.30323E-06 | 1.95452E-06 | 1 | NO | NEGATIVE |
| XACN5 | 6.98434E-06 | 1.71416E-05 | 5 | NO | NEGATIVE |
| XACN6 | 2.35104E-05 | 2.47682E-05 | 1 | NO | NEGATIVE |
| XACN9 | 4.19883E-06 | 1.44199E-05 | 2 | NO | NEGATIVE |

### Immunofluorescence staining of FFPE slides

For these experiments, 6 TLS^+^ and 6 TLS^-^ cervical tumor FFPE slides were selected from the 34 total samples. Selected samples had to exhibit the highest B cell and DC density scores in TLS^+^ tumors and the lowest scores in TLS^-^ tumors. To note, these samples were also analyzed by mass cytometry (CyTOF) using the myeloid panel the lymphoid panel or both.

All reagents and antibodies used for multiplex IF assay are listed in **Supplementary Tables S1–3**. Epitope retrieval was performed with the Target Retrieval solution at pH 6.0 or pH 9.0, depending on the primary antibody tested. Slides were incubated in a water bath for 30 min at 97°C. After washing with TBS, tissue sections were treated with 3% H_2_0_2_ at RT for 30 min, washed again and incubated with Protein Block solution for 30 min at RT. To visualize TLS, tissue sections were then incubated with anti-DC-LAMP antibody for 1 hour at RT. After washing with TBS-0.04% Tween 20, tissue sections were incubated with goat IgG anti-rat IgG1-HRP for 30 min at RT. After washing the tissue sections with TBS-0.04% Tween-20, a tyramide conjugated to AF594 was incubated for 10 min at RT. After washing, the tissue sections were incubated with the Target Retrieval solution for 10 min at 97°C. This step allowed removal of the bound primary and secondary antibodies while protecting AF594 labeling of the DC-LAMP^+^ cells. It also enabled subsequent labeling of the same tissue sections with another antibody. The other antibodies were tested after DC-LAMP labeling in the following order: anti-CD3 revealed with F(ab)’_2_ donkey anti-rabbit IgG-HRP and tyramide-CF514; anti-PanCk revealed with goat anti-mouse IgG1-HRP and tyramide-AF488; anti-CD21 revealed with goat anti-mouse IgG1-HRP and tyramide-AF555; anti-CD20 revealed with goat anti-mouse IgG2a-BV480; and anti-PNAd revealed with F(ab)’_2_ goat anti-rat IgGM-AF647.

To study the B cell, CD8^+^ T cell, and myeloid compartments, three other IF panels were performed using the same techniques and various antibody combinations: anti-CD1a revealed with goat IgG anti-mouse IgG1-HRP and tyramide-CF514; anti-CD8 revealed with goat IgG anti-mouse IgG1-HRP, tyramide-biotine and streptavidine conjugated to AF647; anti-CD68 revealed with goat IgG anti-mouse IgG1-HRP and tyramide-CF430; anti-CD138 revealed with goat anti-mouse IgG1-HRP and tyramide-CF430; anti-CD163 revealed with goat anti-mouse IgG1-HRP and tyramide-AF594.

Nuclei were then stained with DAPI for 5 min at RT before coverslips were mounted onto the glass slides with Fluorescent Mounting Medium. Slide imaging was performed using a Zeiss Axio Observer Z1 at 385 nm, 430 nm, 475 nm, 511 nm, 555 nm, 590 nm and 630 nm with the corresponding filters and Zen software (Zeiss).

### Spatial analysis of cervical tumor FFPE sections

TLS quantification and density analysis were performed using HALO® software (Indica Labs). ROI were delineated, and Hematies, Blank and Necrosis regions were excluded. A classifier was used to discriminate TLS, tumor nest and stroma via staining of PanCk, CD20 and CD3. A Nuclei Segmentation Artificial Intelligence algorithm allowed identification of cells based on their nuclei staining by DAPI (for cell segmentation please see **Supp. Figure 1B**). A Multiplex IHC v3.2 analyzer was used to count CD20^+^ and DC-LAMP^+^ cells in immunohistochemistry slides. A HighPlex FL v4.2 analyzer was used to count cells in IF slides. Spatial analyses with proximity analysis setup were used to measure the distance between immune cell types.

### Tumor sample dissociation

Tumor and healthy cervix samples were processed within 4 hours following surgery. Samples were minced and then dissociated using a gentleMACS dissociator (Miltenyi Biotech) in RPMI 1640 culture medium. Fragments were centrifuged and supernatants were frozen for soluble factor dosage. Fragments were then suspended in Gentle Dissociation Solution (STEMCELL Technologies) and incubated for 20 min at 37°C and 5% C0_2_. Samples were again dissociated using the gentleMACS dissociator before centrifugation and filtering through a 100 µm strainer. Finally, cells were counted and frozen in heat-inactivated fetal calf serum 10% DMSO until staining.

### Soluble factor dosage

#### Chemokines, cytokines, and growth factors

Supernatants were thawed at RT and used for growth factor, cytokines, and chemokines dosage by Luminex using the 48-Plex pan-human cytokine kit (BioRad) according to the manufacturer procedure. Briefly, supernatants were incubated with analyte-specific antibody-coated microbeads for 2 hours. Supernatants were then washed and incubated with a PE labeled secondary antibody for MFI detection. Data were acquired on a BioPlex-200 system (BioRad) and analyzed on BioPlex Manager 6.0 (BioRad), comparing against standard curves for each analyte and according to manufacturer instructions. Raw data were normalized against tumor sample weights (in grams).

#### Immunoglobulins

Supernatants were thawed at RT and used for IgG1-2-3-4, IgA, IgE, IgM and IgD dosage by LEGENDplex (8 plex immunoglobulin isotyping panel, BioLegend) according to the manufacturer procedure. Briefly, supernatants were incubated with analyte-specific antibody-coated microbeads for 2 hours. Supernatants were then washed and incubated with a PE-labeled secondary antibody for MFI detection. Data were acquired on a LSRII (Becton-Dickinson) and Diva software and analyzed on LEGENDplex data analysis software, comparing against standard curves for each analyte. Raw data were normalized against tumor sample weights (in grams).

### Flow cytometry staining and acquisition

Mononuclear cells were stained for live/dead Aqua, CD45-FITC, CD5-APC, CD19-PE-Cy7, CD20-PE-Cf594, CD86-BV645, CD69-PE-Cy5.5, CD38-BV605, IgD-V450, CD27-BV785, CD21-BV711 and CD24-PE for 30 min at 4°C in PBS containing 2% FCS. Cells were then washed with PBS and fixed in 4% PFA for 20 min at 4°C. After PBS washes, data were acquired on a LSRII (Becton-Dickinson). Marker expression was compared against isotypic controls.

### Mass cytometry staining and acquisition

Tumor cells were incubated with cisplatin (1 µM) to stain dead cells. Non-specific epitope binding was blocked with 0.5 mg/mL human Fc Block (BD Biosciences, San Jose, CA, USA). Mononuclear cells were stained for extracellular epitopes for 1h at 4°C, followed by permeabilization (FoxP3 permeabilization reagent, eBioscience, San Diego, CA, USA) for 30 min at 4°C and intracellular marker staining with the panels described in **Suppl. Tables 4–5**. Cells were then washed and labeled overnight with 125 nM iridium intercalator (Fluidigm, South San Francisco, CA, USA) in 2% PFA (Thermo-Fisher). Finally, cell pellets were suspended in Milli-Q water (Merck Millipore, Burlington, MA, USA) containing 10% EQ four element calibration beads (Fluidigm) and filtered through a 35 μm membrane before acquisition on a mass cytometer (Helios® instrument, Fluidigm). Mass Cytometry allows absolute quantification of cells by normalization against EQ™ Four Element Calibration Beads. The bead counts are known as well as the mean Di of all the masses. Beads are identified by 140Ce which is not used for antibody labelling. The normalization factor is the ratio of median Di values to EQ bead population median Di values of the encoding isotopes which cover an extensive portion of the mass range measurable on the Helios instrument. The normalization factors for mass channels between the encoding isotopes are linearly interpolated. All mass channel values for all events are then multiplied by these normalization factors to obtain the normalized values, and data is written to the normalized file. Absolute counts were normalized by weight of tumor (in grams) before comparison between samples.

### Cytometry High-dimensional analysis

#### Flow cytometry analysis

B cell subsets were manually gated based on the expression of CD5, IgD, CD38, CD86, CD21, CD27, and CD24. This allowed the identification of Germinal center (GC) B cells, Memory B cells, Naïve B cells, Plasmablasts and pre-GC B cells. Memory B cells were also subdivided into activated memory, resting memory, intermediate memory and tissue like memory B cells. Subsets were quantified using FlowJo and statistical significance was assessed using one-way ANOVA, Kruskal Wallis test with Dunn’s multiple comparison (GraphPad Prism). Absolute counts were normalized by weight of tumor (in grams) before comparison between samples.

#### Mass cytometry analysis

Myeloid cells (CD45^+^CD3^-^CD19^-^CD56^-^CD33^+^) and T cells (CD45^+^CD3^+^CD19^-^CD33^-^) were manually gated, then exported using FlowJo (Becton-Dickinson). Data were arcsinh-transformed with a cofactor of 5. Myeloid and T cell populations were automatically defined using h-SNE, which allows a better clustering of rare populations such as tumor-infiltrating cells (using default settings of 30 perplexity and 1,000 iterations, Cytosplore V2.2.1)(21,22), and density clustering was analyzed. For myeloid cells, CD11b, CD1c, CD1a, CD11c, HLA-DR, CD123, CD303, CD141, CD163, CD206, PD-L1, CD40, IL-6, and CD209 were used for clustering. For T cells, CD4, CD8, CD25, CD127, CTLA-4, PD-1, ICOS, CD69, CCR7, CD45RA, and TIM3 were used for clustering.

### TLS associated myeloid cell population prediction

Myeloid immune cell signature associated with TLS was created using Least Absolute Shrinkage and Selection Operator (LASSO) regression to avoid overfitting due to a large number of parameters with a limited number of patients (23). Myeloid cell populations that could predict TLS status were identified among the 101 myeloid cell populations described by mass cytometry. Model was regressed on those variables, with penalization and feature selection. Selected variables were then plotted according to TLS status for representation. **Suppl. Table 6** shows the input parameters for the LASSO analysis. LASSO was performed using RStudio V1.3.1093, R-4.0.3.

### Pagès Multiple factor analysis (MFA)

To compare both clinical (FIGO 2018 classification, Age, LN, Emboli, Tumor size) and biological (DC count, B cell count, DC density, B cell density, TLS) parameters we performed a Pagès Multiple factor analysis (MFA), which determines the contribution of each parameter to the dimensions of a principal component analysis. In our case it allowed to determine whether TLS and associated immune factors were associated with negative parameters such as higher FIGO stage or lymphnode status or not. MFA allows the comparison of both quantitative (DC count, B cell count, Age) and qualitative parameters (LN, TLS, DC density, B cell density, FIGO, Tumor size, Emboli). FIGO I and II were grouped as a single variable since only few stages I were available in our cohort. Hence, 3 groups were used for FIGO staging, FIGO low (stages I-II, localized tumors), FIGO intermediate (int., stage III, LN invasion) and FIGO High (stage IV, LN invasion and metastasis). Histology was separated in 2 groups, SCC and others (2 ADK and 1 Mixed histology). B cell and DC densities were scored using quartiles, as we did for TLS maturation scoring. MFA was performed using RStudio V1.3.1093, R-4.0.3. R scripts performing MFA are provided in **Suppl. File S1** and input data are available in **Suppl. Table 7**.

### Multivariate analyses

A multivariate analysis was performed using the following parameters on the n=34 patient samples, TLS^+^ and TLS^-^, FIGO stages low, int and high (similarly to those used for the MFA) and Histology SCC and Others. These 3 parameters (7 variables) allow to statistically discriminated elements of better or poorer prognosis, knowing that increased FIGO stages are of poor prognosis. Similarly, a multivariate analysis was performed on the TCGA dataset using survival parameters, TLS-signature enrichment from 2 different articles (Goc + Coppola), tumor stages, lymphnode (LN) invasion, age and immune cell population obtained from CIBERSORTx enrichment. Multivariate analysis was performed using RStudio V1.3.1093.

### Cell enrichment analysis

CIBERSORTx (Stanford University) was used for the immune cell enrichment analysis of the CESC TCGA database. The individual gene expression file was uploaded to CIBERSORTx (24), and run using both the relative and absolute modes, signature: LM22, 500 permutations, and disabled quantile normalization (recommended for RNA-Seq data). For TLS scoring, enrichment scores for B cells, aDCs, and CD8^+^ T cells were scored as 1 (unenriched), 2 (poorly enriched), or 3 (highly enriched) by tiertile. Noteworthy, the minimun enrichment score value considered usable by CIBERSORTx is 0.02. Samples with high enrichment in every cell subset were considered TLS^+^. Results from CIBERSORTx enrichment are available in **Suppl. Table 8**.

### Statistical analysis

TLS^-^ and TLS^+^ tumors were discriminated by the morphology of T + B cell aggregates, their presence *in situ*, and the quartile classification of B cell densities (1^st^ quartile = 3.37 × 10^-6^ cells/µm², median = 3.31×10^-5^ cells/µm², 3^rd^ quartile = 1.67.85 × 10^-4^ cells/µm²) and aDC densities (1^st^ quartile = 70.19 × 10^-6^ cells/µm², median = 2.15×10^-5^ cells/µm², 3^rd^ quartile = 4.85 × 10^-5^ cells/µm²). The 1^st^ quartile shows the lowest and the 4^th^ quartile the highest densities. Aggregate presence and densities were then scored according to the above-described parameters and tumors labeled as TLS^-^ and TLS^imm.^ and TLS^mat.^ (**Supp. Fig. 1**).

Statistical analyses were generated using GraphPad Prism V8.00, with data expressed as mean ± standard error of the mean. Statistical significance between two groups was calculated using the non-parametric Mann–Whitney test, while multiple group comparisons were calculated using the non-parametric Kruskal-Wallis test followed by a Dunn, multiple comparison post-test. A *p*-value < 0.05 was considered as significant. For survival analyses, PFS was defined as the time from diagnosis until death or relapse, whichever occurred first. Patients without an event were censored at the time of their last follow up. Survival times were estimated by the Kaplan–Meier method and compared using the log-rank test.

## Results

### TLS maturation levels vary in cervical tumors and is associated with soluble microenvironment alterations

Cervical tumor samples (n=34) were screened for the presence of TLS according to CD20 (B cells), CD21 (follicular dendritic cells, fDCs), CD3 (T cells) and DC-LAMP (activated DCs, aDCs) (**Fig. 1A**). First, B cell aggregate presence were assessed by CD20 staining (**Supp. Fig 1A.**). B cell aggregates were then associated with the presence of T cells (CD3^+^) and aDCs (DC-LAMP^+^). B cell and aDC density and the presence of fDCs were then used as TLS maturation parameters. This allowed to divide our cohort into TLS^-^ (no aggregates), TLS^lo^ (presence of lymphoid aggregates with low B cell and aDC density), immature TLS (TLS^imm.^, higher B cell and aDC density) and mature TLS (TLS^mat.^, highest B cell and aDC density with fDC presence) (**Fig. 1B, C and Supp.Fig 1A**.). We found TLS^imm.^ or TLS^mat.^ presence in 50% of our cohort samples. TLS maturation stages were validated against manual TLS counts and showed increased TLS counts in TLS^imm.^ and TLS^mat.^ samples (**Fig. 1D**). As B cell and aDC density did not vary between TLS^-^ and TLS^lo^ samples, they were analyzed as a single group in the rest of the study and called TLS^-^. **Fig. 1E** displays a mature TLS exhibiting a B cell zone (CD20^+^ B cell and CD21^+^ FDC cluster) and a T cell zone (CD3^+^ T cell and DC-LAMP^+^ aDC cluster) and the presence of high endothelial venules (PNAd^+^ cells). Next, we measured secreted cytokines, soluble effectors and chemokines in TLS^mat.^, TLS^imm.^, TLS^-^ tumor and healthy cervix dissociation supernatants by Luminex. CXCL1 was only significantly higher in TLS^mat.^ samples compared to healthy donors. IL-2ra, IL-16, CCL3, CXCL9 and CXCL10 were increased in TLS^mat.^, TLS^imm.^ and TLS^-^ tumor supernatants compared to healthy cervix supernatants. CCL4, was increased in TLS^mat.^ and TLS^imm.^ tumor supernatants compared to healthy cervix supernatants. TNF-α, TRAIL and CCL5 were increased in TLS^imm.^ and TLS^-^ tumor supernatants compared to healthy cervix supernatants, but not in the TLS^mat.^ supernatants. Overall, no significant differences were observed between TLS^mat.^, TLS^imm.^ and TLS^-^ conditions (**Fig. 1F**). In summary, our results show that TLS are present in 50% of cervical tumors in our cohort, are associated with higher densities of B cells and aDC, and tends to be associated with increased CXCL1.

**Figure 1:**
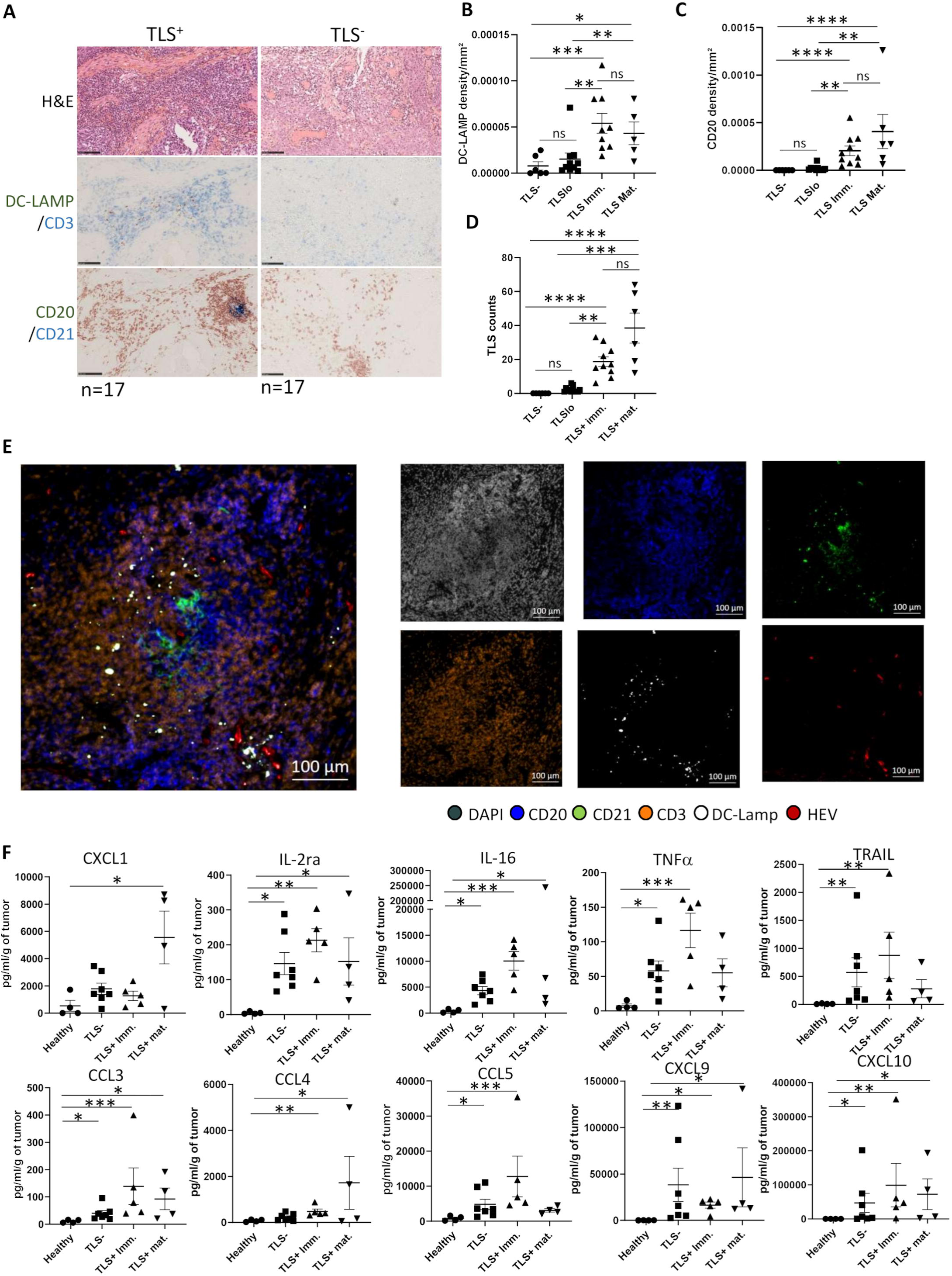
TLS are associated with B cell density and dendritic cell activation as well as with a higher inflammatory profile in cervical tumors. **A**, Immunohistochemistry (IHC) for DC-LAMP, CD3, CD20 and CD21 was performed on n=34 cervical tumor FFPE samples. TLS were detected in 17/34 samples. **B**, Dot plots show DC density against the TLS maturation classification **C**, Dot plots show B cell density against the TLS maturation classification **D**, Dot plots show TLS counts against the TLS maturation classification in the 34 FFPE samples. **E**, Images show a representative TLS, stained by multiplex IHC for CD20, CD3, DAPI, DC-LAMP, CD21 and PNAD. **F**, soluble factors were detected in tumor supernatants by Luminex. Dot plot represent the concentration of each proteins in healthy cervix, TLS^-^, TLS^imm.^ and TLS^mat.^ tumors in pg/ml. Statistical analyses were performed using Kruskal-Wallis test followed by a Dunn, multiple comparison post-test (*p-val<0.05, **p-val<0.01, *** p-val<0.005, **** p-val<0.001).

### TLS^+^ tumors exhibit a higher B cell infiltrate characterized by an enrichment of plasmablasts in the stromal compartment

We analyzed the B cell compartment in TLS^+^ and TLS^-^ tumors by flow cytometry and unsupervised analysis. B cell subsets were identified by manual gating according to the expression of CD19, CD20, CD5, IgD, CD27, CD24, CD38 and CD21 (**Fig. 2A**). The absolute number of B cells, normalized by grams of tumor sample, was higher in TLS^+^ tumors as compared to TLS^-^ tumors (**Fig. 2B**). Germinal center B cells (GC, CD38^+^IgD^-^), pre-GC (CD38^+^IgD^+^), Memory B cells (CD38^-^IgD^-^), naïve B cells (CD38^-^IgD^+^) or plasmablasts (CD38^hi^IgD^-^) did not vary significantly between TLS^-^ and TLS^+^ tumors (**Fig. 2C**). Among memory B cells, we identified Tissue Like Memory B cells (TLM, CD27^-^CD21^-^), Intermediate memory B cells (CD27^-^CD21^+^), resting memory B cells (CD27^+^CD21^+^) and activated memory B cells (CD27^+^CD21^-^). None of these subset frequencies were significantly different in TLS^+^ vs. TLS^-^ tumors (**Fig. 2D**). Hence, we investigated the spatial localization of B cells within the cervical tumor and stroma by multiplex immunofluorescence (IF) (**Fig. 2D)**. Briefly, we applied the tumor nest region of interest (ROI), determined as pan-cytokeratin positive (PanCk^+^), to 6 TLS^+^ and 6 TLS^-^ samples. The absence of PanCk staining identified the stroma ROI. Also, all immune cells were PanCk^-^. We observed no significant differences in CD20^+^CD138^-^ B cells, CD20^+^CD138^+^ plasmablasts, and CD20^-^CD138^+^ plasma cells between TLS^-^ and TLS^+^ samples in the tumor nest (**Fig. 2E**). While CD20^-^ plasma cells and B cells infiltrate did not vary, plasmablasts infiltrated TLS^+^ stroma significantly more than TLS^-^ stroma (**Fig. 2F**). As plasmablasts increased in TLS^+^ stroma, we assessed soluble immunoglobulins (Ig) concentration in TLS^mat.^, TLS^imm.^, TLS^-^ and healthy cervix supernatants, and found significant increases in IgG1, IgG2, IgG3, IgA, IgM and IgD in TLS^mat.^, TLS^imm.^, TLS^-^ sample supernatants compared to those of healthy cervix (**Fig. 2G**). Here, we observed no statistically significant differences between the TLS^-^, TLS^mat.^ and TLS^imm.^ Samples. IgG4 was solely significantly increased in TLS^mat.^ tumors compared to healthy cervix. Interestingly, TLS^mat.^ showed increased levels of IgE compared to healthy cervix, TLS^-^ samples and a close to significant increase compared to TLS^imm.^ supernatants (*p*-val = 0.061). (**Fig. 2G**). Altogether, these results suggest enrichment of the B cell compartment marked by enriched plasmablasts in TLS^+^ tumors which may increases the secretion of IgE.

**Figure 2:**
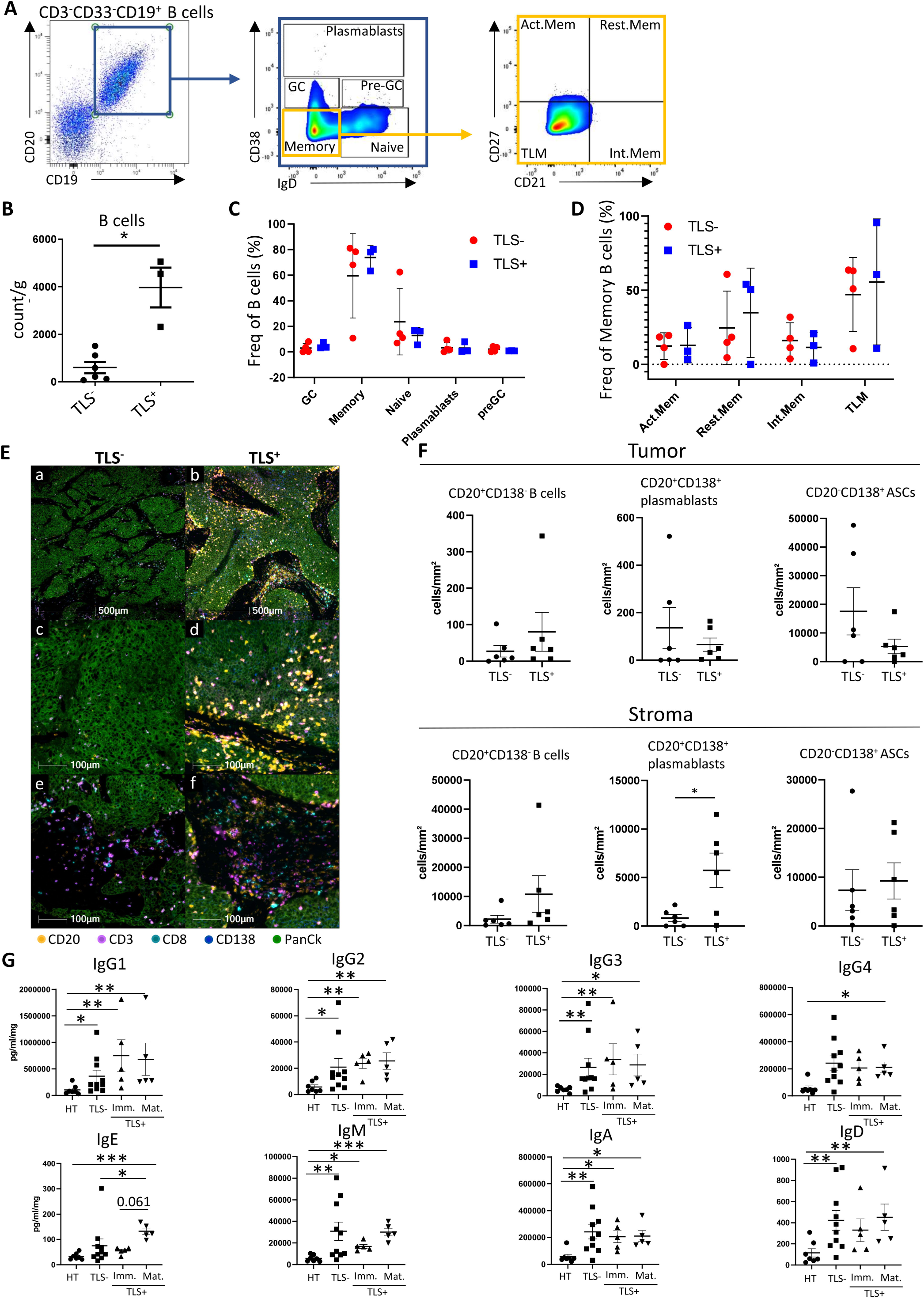
TLS^+^ tumors exhibit a higher B cell infiltrate characterized by an enrichment of plasmablasts in the stromal compartment and exacerbated IgG presence. **A,** Manual gating of tumor infiltrating B cell subpopulations in n= 6 TLS^-^ and n=3 TLS^+^ samples. **B**, B cell absolute count normalized by gram of tumor sample. **C**, B cell subpopulation frequencies in n= 6 TLS^-^ and n=3 TLS^+^ samples. **D**, Memory B cell phenotyping in n= 6 TLS^-^ and n=3 TLS^+^ samples. Frequencies of parent population are shown only when parent population contained enough events (>30 cells). **E**, Multiplex IF was performed for CD20, CD138, CD3, CD8, and PanCk on TLS^-^ (left, n=6) and TLS^+^ (right, n=6) FFPE samples. **F**, B cells (CD20^+^CD138^-^) and ASCs (CD138^+^CD20^+^, CD138^+^CD20^-^PanCk^-^) were quantified in the tumor (PanCk^+^) compartment. **G**, B cells (CD20^+^CD138^-^) and ASCs (CD138^+^CD20^+^, CD138^+^CD20^-^PanCk^-^) were quantified in the stromal (PanCk^-^) compartment. **H**, immunoglubulins were measured in healthy cervix, TLS^-^, TLS^imm.^ and TLS^mat.^ supernatants. Statistical analyses were performed using Kruskal-Wallis test followed by a Dunn, multiple comparison post-test (\**p*-val<0.05, \*\**p*-val<0.01, *** *p*-val<0.005).

### CD163^+^ macrophages and activated cDC2 are enriched in TLS^+^ cervical tumors

The myeloid compartment is known to regulate TLS formation (macrophages)(25,26) and to participate in their T cell-activation function (aDCs)(27). Therefore, we characterized the myeloid compartment by mass cytometry and identified 8 clusters composed of Langerhans cells (LCs, CD1a^+^CD11c^+^), CD1a^lo^ DCs (CD1c^+^CD1a^lo^), cDC1 (HLA-DR^+^CD11c^+^CD11b^-^CD141^+^), cDC2 (CD11c^+^CD1c^+^), pDCs (CD123^+^CD303^+^), early myeloid-derived suppressor cells (eMDSCs, HLA-DR^-^/^lo^CD11b^+^CD14^-^CD15^-^), monocytic MDSCs (mMDSCs, HLA-DR^-/lo^CD11b^+^CD14^+^) and TAMs (CD64^+^CD68^+^HLA-DR^+^) (**Suppl. Fig. 2A, 1B**). We also divided TAMs according to CD163 expression (**Fig. 3A; Suppl. Fig. 2A**). TAM numbers did not vary significantly between TLS^-^ and TLS^+^ tumors. However, the frequency of CD163^+^ TAMs was higher in TLS^+^ tumors compared to TLS^-^ tumors (**Fig. 3B**). *In situ* analysis of TAMs showed a trend for increased CD163^+^ TAM infiltrate in TLS^+^ sample stroma and tumor (**Fig. 3C**). Next, we assessed the infiltration of TLS^+^ and TLS^-^ samples by cDC2, LCs and CD1a^lo^ DCs, and observed an increased number of cDC2 in TLS^+^ samples compared to TLS^-^ samples (**Fig. 3D**). Therefore, we investigated cDC2 localization using the DC-LAMP marker for aDCs and CD1a for LCs. We found no difference in aDCs (CD1a^-^DC-LAMP^+^), activated LCs (aLCs, DC-LAMP^+^CD1a^+^), and immature LCs (iLCs, DC-LAMP^-^CD1a^+^) infiltrate in the tumor ROI (PanCk^+^). However, aDCs infiltrate was significantly higher in the TLS^+^ sample stroma (**Fig. 3E**). In summary, our results show an increase in CD163^+^ macrophages and aDCs in the stroma of TLS^+^ samples.

**Figure 3:**
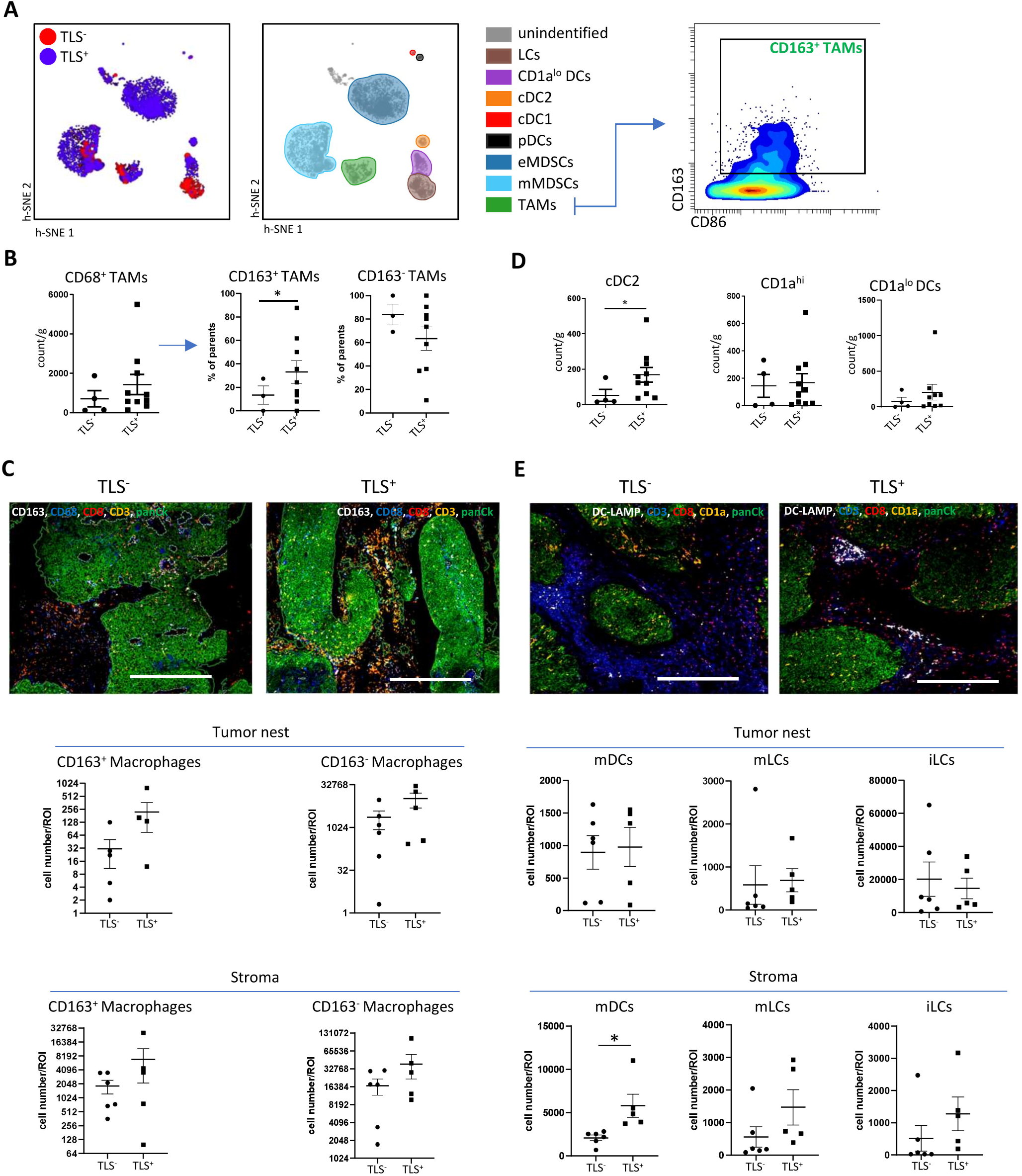
CD163^+^ macrophages and mature cDC2 are enriched in TLS^+^ cervical tumors. **A**, Tumor infiltrating myeloid cells were analyzed by mass cytometry on n= 14 samples, h-SNE and density clustering. Clusters were annotated according to the expression of markers in **Supplementary Table 5**. **B**, TAMs were gated and quantified in n=14 samples and plotted according to the presence or absence of TLS. **C**, multiplex IHC was performed to identify CD163^+^CD68^+^ and CD163^-^CD68^+^ TAMs in TLS^+^ (n=6) and TLS^-^ (n=6) tumors. Representative images are shown (top) and cell quantities were assessed (bottom). **D**, cDC2, CD1a^hi^ LCs and CD1a^lo^ DCs were gated and quantified in n=14 samples and plotted according to the presence or absence of TLS. **E**, multiplex IHC was performed to identify mDCs (DC-LAMP^+^CD1a^-^), mLCs (DC-LAMP^+^CD1a^+^) and iLCs (DC-LAMP^-^CD1a^+^) in TLS^+^ and TLS^-^ tumors. Representative images are shown (top) and cell quantities were assessed in the tumor nest (center) and stroma (bottom). Statistical analyses were performed using Mann-Whitney test (\**p*-val<0.05)

### Myeloid cells are more activated and better differentiated in TLS^+^ tumors

We investigated the phenotypic variations within myeloid cell subsets. We assessed the expression of PD-L1, CD40, LILRB2, CD1d, CCR2, CCR5, CXCR4, CD80, CD86, HLA-DR, PVR, Nectin2, IL-6, CD10, CD206, CD209 and CD141 among LCs, CD1a^lo^DCs, cDC1, cDC2, pDCs, eMDSCs, mMDSCs, CD163^-^ TAMs, and CD163^+^ TAMs. The frequencies were filtered on a 10% average expression and a minimum of 30 cells per cluster. We analyzed the resulting 101 variables (44 TLS^+^, 57 TLS^-^) using the LASSO algorithm to obtain each variable predictive of either a TLS^+^ or TLS^-^ profile (**Fig. 4A**). CD163^+^ TAMs were the most highly represented cell type in the TLS^+^ group, with six of these variables corresponding to the contribution of CD1d, CXCR4, PD-L1 and CCR5 (**Fig. 4B**; **Suppl. Table 6**). CD40 expression, which marks activation, on cDC2 was also associated with the TLS^+^ environment (**Fig. 4B**), which is logical considering that the presence of aDCs is one of the main features of a TLS^+^ microenvironment. Early MDSCs contributed to the TLS^+^ group with the expression of CCR2 (**Fig. 4C**). IL-6 expression on CD163^-^ TAMs and CD1a^lo^ DCs were also selected by the LASSO analysis (**Fig. 4C**). PVR^+^CD1a^lo^ DCs (poliovirus receptor, TIGIT ligand) was also selected by the LASSO algorithm. Finally, the presence of mMDSCs associated with a TLS^-^ microenvironment (**Fig. 4C**). These findings were validated by a univariate analysis conducted as a post-test. All variables associated with a TLS^+^ microenvironment were significant, while only IL-6^+^CD163^-^ TAMs were significant in a TLS^-^ microenvironment (Log2FC>1 and *p*-val<0.05) (**Fig. 4D)**. Overall, the myeloid cell contribution to a TLS^+^ environment is determined by the expression of activation (CD40) and inhibition (PD-L1) markers, which reflects a better differentiation of TAMs and cDCs. Furthermore, the contribution of chemokine receptor expression (CXCR4, and CCR5) by TAMs also indicates an increase in the recruitment of these cells. Finally, our results show the importance of the myeloid cell contribution to a TLS^+^ environment, which reflects a greater differentiation and activation of mainly macrophages, and DCs.

**Figure 4:**
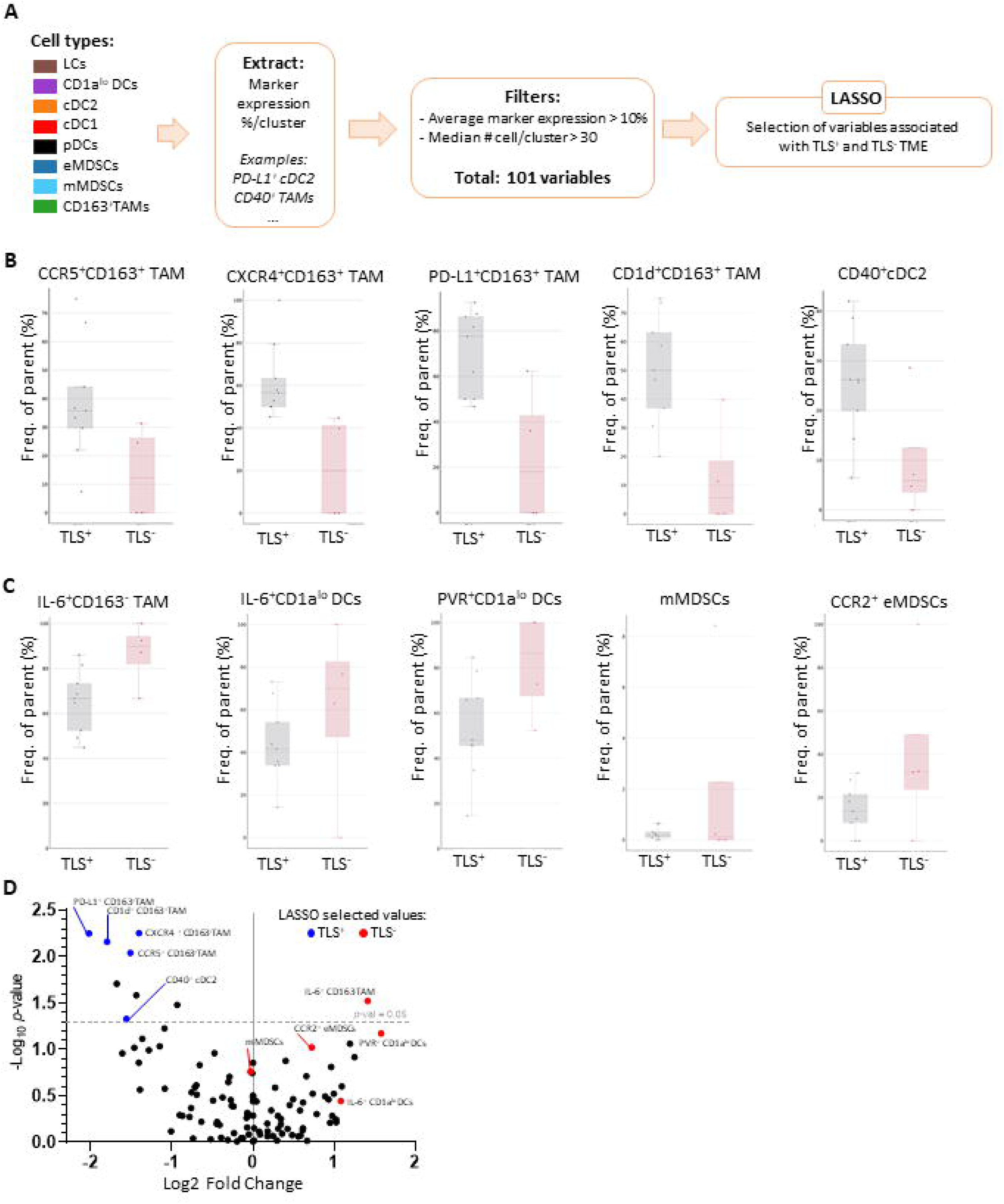
LASSO analysis highlights macrophage and DC activation in TLS^+^ tumors. **A**, Individual marker frequencies (PD-L1, CD40, CD86 …) was assessed in each myeloid cell subtype identified by CyTOF per patient (n=14). Then, data were filtered to retain frequencies in cell cluster with a count > to 30 cells as well as an average expression > to 10%. This led to 101 identified variables, which were then analyzed by LASSO to identify specific variables to either TLS^-^ or TLS^+^ TME. **B**, LASSO selected variables associated with TLS^+^ TME. **C**, LASSO identified variables associated with TLS^-^ TME. **D**, univariate analysis of the 101 variables. Volcano plot shows distribution of variables in TLS^-^ and TLS^+^ TME.

### Spatial analysis reveals increased interactions between aDCs and CD8^+^ T cells, which exhibit higher diversity in TLS^+^ tumors

We hypothesized that the involvement of aDCs in TLS^+^ tumors was associated with their proximity to CD8^+^ T cells, and could enhance immune response. Therefore, we performed cell segmentation and identified aDCs (DC-LAMP^+^CD1a^-^), activated CD1a^+^ DCs (aCD1a^+^ DC, DC-LAMP^+^CD1a^+^), and CD8^+^ T cells **(Fig. 5A)**. Next, we quantified the number of CD8^+^ T cells within a 10 µm, 10–20 µm and 20–30 µm range of either a single aDC or aCD1a^+^ DC (**Fig. 5B**). The number of CD8^+^ T cells within a 10 µm, a 10–20 µm and a 20–30 µm range of an aCD1a^+^ DC tended to increase in TLS^+^ compared to TLS^-^ tumors, although the difference was not significant (**Fig. 5C**). However, the number of CD8^+^ T cells within a 10 µm range of an aDC significantly increased in TLS^+^ tumors compared to TLS^-^ tumors (**Fig. 5D left**), and remained unchanged in the 10–20 µm and 20–30 µm ranges. This close proximity suggests that aDCs directly interact with CD8^+^ T cells more efficiently in TLS^+^ tumors, which may improve tumor control (**Fig. 5D**).

**Figure 5:**
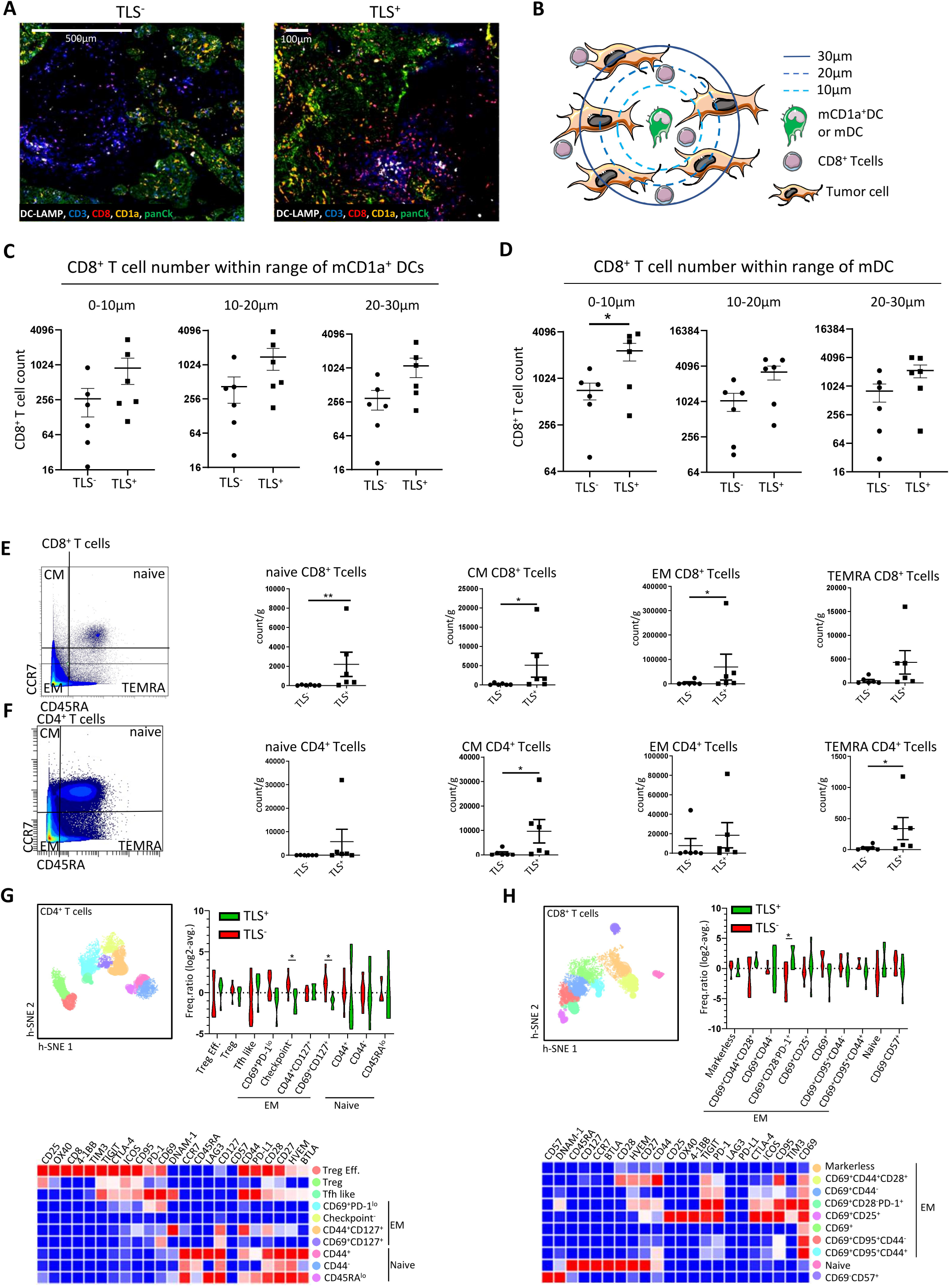
Spatial analysis reveals increased interactions between mDCs and CD8^+^ T cells, which exhibit higher diversity in TLS^+^ tumors. **A**, multiplex IF was performed to identify mDCs (DC-LAMP^+^CD1a^-^), mCD1a^+^ DCs (DC-LAMP^+^CD1a^+^) and nmCD1a^+^ DCs (DC-LAMP^-^CD1a^+^) in TLS^+^ (n=6) and TLS^-^ (n=6) tumors. **B**, schematic representation of proximity analysis. **C**, mCD1a^hi^ proximity to CD8^+^ T cells was quantified within 10µm, 10-20µm and 20-30µm range in TLS^-^ and TLS^+^ tumors. **D**, mDCs proximity to CD8^+^ T cells was quantified within 10µm, 10-20µm and 20-30µm range in TLS^-^ and TLS^+^ tumors. **E**, CD8^+^ T cells subsets (naïve, EM, CM, TEMRA) were gated and quantified by mass cytometry (n=14). Absolute counts are shown as dot-plots. **F**, CD4^+^ T cells subsets (naïve, EM, CM, TEMRA) were gated and quantified by mass cytometry. Absolute counts are shown as dot-plots. **G**, CD4^+^ T cell analysis by h-SNE and density clustering. The heatmap represent maker expression per cluster and each cluster annotation. Violin plot represent the frequency of each cluster in TLS^+^ and TLS^-^ samples. “Checkpoint^-^“ EM CD4^+^T cells did not express any immune checkpoint present in our panel. **F**, CD8^+^ T cell analysis by h-SNE and density clustering. The heatmap represent maker expression per cluster and each cluster annotation. Violin plot represent the frequency of each cluster in TLS^+^ and TLS^-^ samples. “Markerless” CD8^+^ T cell did not express any of the markers present in our Lymphocyte mass cytometry panel other dans CD3 and CD8. Statistical analyses were performed using Mann-Whitney test (\**p*-val<0.05).

To further dissect the T cell response between TLS^+^ and TLS^-^ tumors, T cell subsets were mapped. We showed that absolute numbers of naïve (CCR7^+^CD45RA^+^), central-memory (CM, CCR7^+^CD45RA) and effector-memory CD8^+^ T cells (EM, CCR7^-^CD45RA^-^) were increased in TLS^+^ tumors, which was not the case for effector-memory re-expressing CD45RA (TEMRA, CCR7^-^CD45RA^+^) CD8^+^ T cells (**Fig. 5E**). The absolute numbers of CD4^+^ T cell subsets also varied between TLS^+^ and TLS^-^ tumors. Indeed, naïve and TEMRA CD4^+^ T cell numbers were increased in TLS^+^ tumors (**Fig. 5F**). Therefore, these results show a greater diversity and differentiation of the T cells and T cell response in TLS^+^ tumors.

Using h-SNE clustering and density clustering, we identified variations due to specific marker and quantified the cluster frequencies (**Fig. 5G, 5H**). Among CD4^+^ T cells, only EM CD4^+^ T cells varied. Indeed, EM CD4^+^ T cells which did not express any specific checkpoint marker (**Fig. 5G** “Checkpoint^-^”) and CD69^+^CD127^+^ EM CD4^+^ T cells were enriched in TLS-tumors compared to TLS^+^ tumors (**Fig. 5G**). Among CD8^+^ T cells, one CD69^+^CD28^-^PD-1^+^ EM CD8^+^ T cells were enriched in TLS^+^ tumors (**Fig, 5H**). In summary, these results show increased aDC/T cell crosstalk and T cell activation and immune checkpoint expression in TLS^+^ tumors compared to TLS^-^ tumors.

### Mature TLS improve prognosis in cervical tumors

We hypothesized that the enhanced activation of immune cells within TLS^+^ tumors could allow patient segregation and predict progression-free survival (PFS). Therefore, we performed a Pages-multiple factor analysis (MFA), which allows patient segregation by quantitative parameters (DC count, B cell count, age) and qualitative parameters (FIGO stage, LN invasion, emboli presence, DC and B cell density, TLS maturation, relapse and death, treatment), and assesses the impact of these parameters on patient survival (**Supp. Fig. 3A, B**). The MFA defined two axes, where Dim.1 (Axis 1) corresponded to immune parameters (TLS and immune cell counts) and Dim.2 (Axis 2) corresponded to disease-related variables (Stages, Treatment) (**Supp. Fig. 3A, B**). LN invasion and emboli presence discriminated patient along Dim.2, while DC and B cell number and densities discriminated patient along Dim.1 (**Supp. Fig. 3C**). Age and Histology did not influence patient segregation (**Supp. Fig. 3C**). Here, we showed that patients were well-segregated along Dim.1 by the different TLS maturation status, especially between mature TLS and the two other groups (**Fig. 6, A**). FIGO stages and discriminated patients along Dim.2. Interestingly, relapses and death segregated patients along Dim.1, and patient that relapsed or died corresponded mainly to patients with immature TLS or not presenting TLS (**Fig. 6, A**). A multivariate analysis was performed between presence or absence of TLS, FIGO stages and Histology and showed that high FIGO stages were associated significantly with death and relapse (*p*-val = 0.009) while TLS presence tended to associate to survival (*p*-val = 0.093) (**Suppl. Fig. 4, A**), which confirmed our results from the MFA.

**Figure 6:**
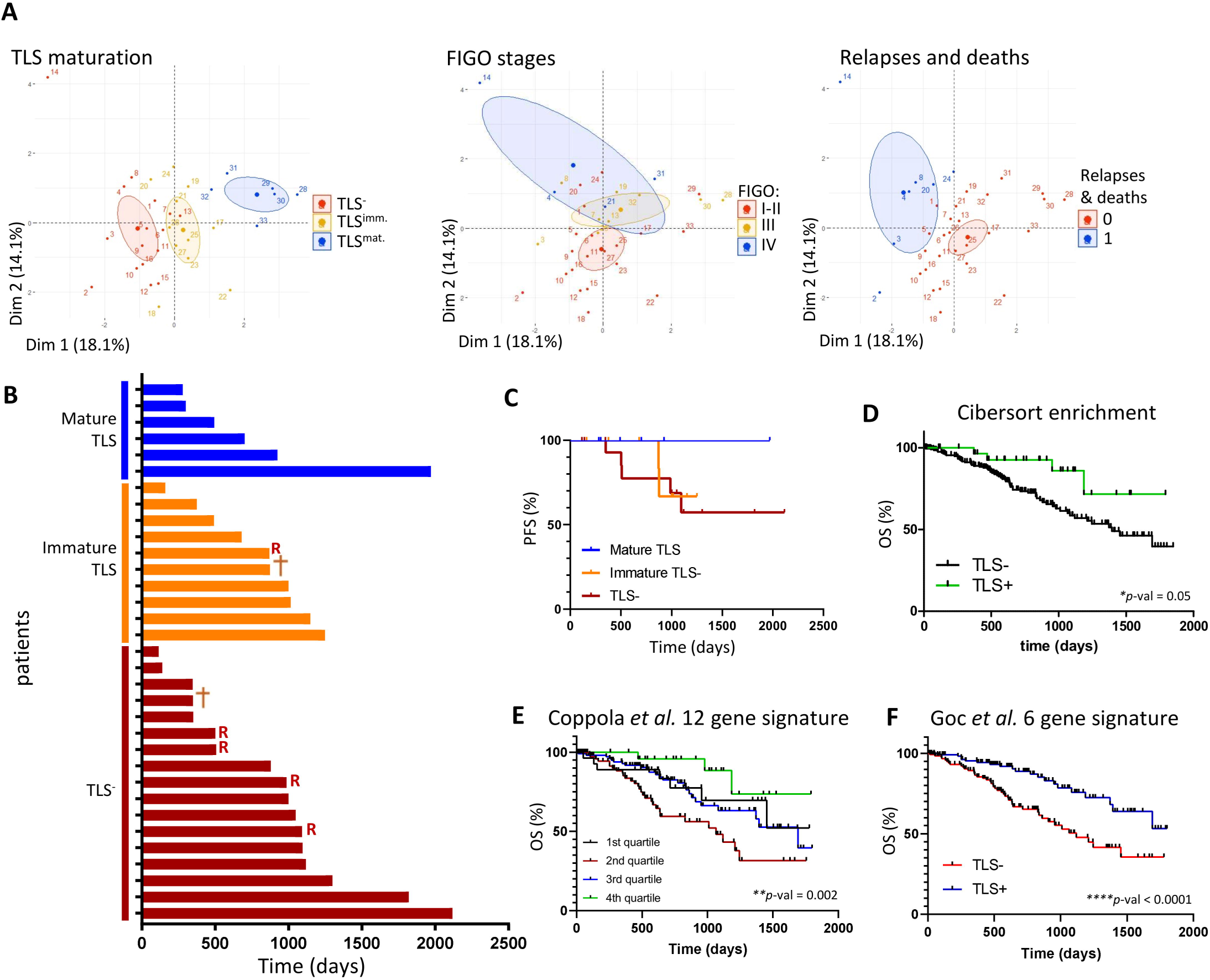
TLS^+^ tumors with high B cell and DC density are associated with improved prognosis. **A.** MFA was performed on 34 patients and highlighted TLS maturation, FIGO stages and Relapses and death as significant discriminating factors. **B,** Waterfall chart displays the follow up of n=34 patients, annotated in function of the absence of TLS (red, TLS^-^), presence of immature TLS^+^ (orange) or mature TLS (blue). Crosses represent deaths and R represent relapses. **C,** Kaplan-Meier plot represent relapse free survival of patients in the 3 groups defined in **B**. **D**, Immune cell enrichment algorithm CIBERSORTx was used to investigate TIL infiltrate in TCGA CESC cohort (306 patients). Kaplan-Meier curves were generated for scored TLS related cell types (CD8^+^ Tcells, B cells and activated DCs). A high score of enrichment for these cell types was annotated TLS^+^ (green) and lower scores were annotated TLS^-^ (black). \**p*<0.05. Coppola et al. (**E**) and Goc et al. (**F**) gene signatures were applied to the CESC dataset from TCGA. The signatures were then scored using quartiles (**E**) or median (**F**) and Kaplan-Meier curves were generated. Statistical differences were calculated using the Log-rank test. \**p*-val<0.05, \*\**p*-val<0.01, \*\*\**p*-val<0.005, \*\*\*\**p*-val<0.001.

Next, we studied patient PFS according to TLS^-^, TLS^imm.^ and TLS^mat.^, and found that the TLS^mat.^ group (n=6 patients) did not include any death or relapse, while the TLS^imm.^ (n=10 patients) included one death and one relapse, and the TLS^-^ group (n=17 patients) included one death and four relapses (**Fig. 6B**). **Fig. 6C** shows better PFS in the TLS^mat.^ group, and similar PFS curves in the TLS^imm.^ and TLS^-^ groups. To validate these results, we performed an immune cell enrichment analysis on the cancer genome atlas CESC dataset (TCGA) using the CIBERSORTx algorithm (**Supp. Table 8**), followed by a survival analysis according to B cell, CD8^+^ T cell, activated DC (aDCs), and CD4^+^ TEM enrichment. B cell and CD8^+^ T cell enrichment significantly improved patient survival, while aDCs only showed a non-significant trend for improved survival and CD4^+^ TEM enrichment did not influence survival outcome (**Supp. Fig. 3D**). Finally, we scored the enrichment for TLS presence, where TLS^+^ tumors are enriched for B cells, aDCs and CD8^+^ T cells and confirmed TLS^+^ tumors association with better survival (**Fig. 6D**). To strengthen these results, we also performed survival analyses using two published TLS-related gene signatures (from Coppola et al.(28) **Fig. 6E** and Goc et al. (27) **Fig.6F**), which again showed better prognosis in patient with enriched TLS-related gene signatures. We also performed a multivariate analysis on the TCGA cohort using TLS gene signature enrichment from *Coppola et al.* and *Goc et al.* to avoid redundancy with CIBERSORTx individual population enrichment, tumor stages (sSTAGEs), LN invasion, metastasis presence, and Age. Multivariate analysis returned that plasmocytes, CD8^+^ T cells and M2 macrophages significantly associated with survival, which matched our results of stromal CD138^+^ and CD8^+^ T cell number increase, as well as CD163^+^ macrophages frequency. Expectedly, LN invasion and advanced tumor stages (T4) associated with death (**Suppl. Fig 4,B**). TLS signatures did not associate with survival in this analysis. This may be due to transcriptional signature being redundant with those used by CIBERSORTx enrichment, or critical parameters lacking such as spatial organization and cell densities within TLS. In summary, mature TLS are associated with improved PFS in cervical cancer.

## Discussion

Our study demonstrates that TLS are associated with a more differentiated and activated immune infiltrate in cervical tumors. Indeed, the presence of TLS is associated with better differentiation and activation of macrophages and DCs, which is linked to a better CD8^+^ T cell recruitment and activation. Altogether, these parameters contribute to a better prognosis in patients with TLS^+^ cervical tumors. Although our cohort is of modest size, our results are robust and validated by multiple statistical methods and unsupervised analyses. Furthermore, they integrate quantitative and qualitative methods, which allows to assess the diversity of cervical tumor microenvironment.

Indeed, we used an integrated approach to investigate TLS and the immune infiltrate in cervical tumors. First, we showed an increase in CXCL1 in TLS^mat.^ TMEs specifically. CXCL1 is usually associated with poorer prognosis in cervical tumors (29), however, the presence of mature TLS seems to counterbalance this effect in our cohort. IL-16, TNF-α, IFN-α, CCL5, CXCL9 and CXCL10 are linked to TLS and increase immune infiltrate and better prognosis in a transcriptomic signature(30). These soluble effectors seemed to increase progressively with TLS maturation, but these changes remained not significant, which may be due to the size of our cohort. IL-16 attracts CD4^+^ T cells, but also macrophages and DCs (31,32). CXCL9 and CXCL10 are produced in response to IFN-γ in the TME, participate to T cell recruitment to hyperplastic epidermis via CXCR3 (33), and associate to prognosis improvement in estrogen receptor-negative breast tumors(34), but promotes progression in tongue cancer(35). CCL5 allows T cell, immature DC and monocyte/macrophage recruitment(36,37), but also allows angiogenesis and tumor growth (38), through the recruitment of macrophages (39), which could be prevented by neutralizing the chemokine (40) or its signaling pathway(38).

Consequently, a TLS^+^ microenvironment is associated with enriched immune cell proportions, which is common among cervical cancer and other HPV^+ (19)^ and HPV^-^ tumors(15,17,41,42). As expected, the analysis of the B cell compartment showed an enrichment in overall B cells within TLS^+^ samples. More specifically, we found an increase in CD138^+^ plasma cells within the stromal compartment of these samples. Plasma cells were associated with cytokine production, chemotaxis and carcinogenesis in a single cell RNA seq study of cervical tumors(43). In our study, plasma cell enrichment was associated with an increased abundance of intra-tumoral IgEs and IgGs in TLS^+^ samples. Tumor specific IgEs are currently investigated as therapeutic strategies (44). Their relation with TLS^+^ TME and activated macrophages will be discussed in the adapted paragraph. To our knowledge, no previous studies exist on the intra-tumoral IgE and IgG presence in cervical tumors. In cervical cancer, anti-HPV neutralizing IgGs mark vaccination efficiency preventing HPV-induced malignancies (45). While natural HPV clearance only allows protection against a specific strain of HPV and generates low IgG titers vaccination allows the targeting of multiple HPV strains and are more abundant(46,47). Therefore, the increased presence of intra-tumoral IgGs in TLS^+^ tumors could be linked to a better response to HPV and/or tumor antigens, which would contribute to improved prognosis in TLS^+^ patients.

Myeloid cells contribute to the regulation of immune responses in tumors(48-52). aDC infiltration is usually associated with a better T cell activation and overall better control of tumor progression (53,54). Here, we show an enrichment of CCR2^+^CD40^+^ aDCs in TLS^+^ samples compared to TLS^-^ samples within cervical tumor and stroma. Here, CCR2 indicates an increased recruitment of aDCs from the blood to the TME. In TLS^+^ samples, aDCs accumulated in the stroma where they interact closely with CD8^+^ T cells. Similarly, Ruffin *et al.* demonstrated that cell-cell neighborhoods are composed of myeloid cells, CD8^+^ T cells, B cells, and conventional T cells in HNSCC tumors that exhibited mature GC^+^TLS and had improved prognosis (19). In ovarian tumors, the presence of antigen presenting cell niches where CD11c^+^ myeloid cells interact with CD8^+^ T cells potentiates anti-CD28 therapy and improves prognosis(55). By analyzing CD8^+^ and CD4^+^ T cell phenotypes we observed a more diverse infiltrate. EM, TEMRA, naïve, CM CD4^+^, and CD8^+^ T cells were all enriched in TLS^+^ tumors, while remaining at low levels in TLS^-^ tumors.

TAMs are often associated with a negative outcome. Indeed, CD204^+^, CD206^+^ and CD163^+^ TAMs are associated with decreased prognosis in cervical cancer (48) and many other tumor types (56-59). Here, we report an increase in the expression of immune checkpoints and ligands (PD-L1, CD40) and chemokine receptors (CCR2 and CCR5) in TLS^+^ tumors. CCR2 and CCR5 expression indicates the blood monocyte origin of TAMs in TLS^+^ tumors. TLS are known to improve response to immunotherapies(60,61) which fits with an increased expression of immunotherapy targets, such as CD40, PD-L1, CCR2 and CCR5, on TAMs and aDCs. This has been clearly established for immune checkpoint inhibitors such as anti-PD-1, anti-PD-L1, and anti-CTLA-4(60,61). Recently, PD-L1^+^ TAM were associated with PD-1^+^ T cells, which appeared to improve survival when the ratio is in favor of CD8^+^ T cell infiltrate in cervical tumors (11-13) and when PD-L1^+^ TAMs were forming niches with T cells rather than being in contact with tumor cells in breast tumors (62), which correspond to features of TLS^+^ TMEs and our Cox regression results. Agonistic anti-CD40 mAbs allow DCs to improve anti-tumoral responses (63-66). We observed increased expression of CCR2 and CCR5 on TAMs from TLS^+^ cervical tumors. Targeting these chemokine receptors represent an innovative strategy to limit TAM infiltration. Indeed, CCR2/CCR5 inhibitors are currently being investigated in colorectal and pancreatic cancers (67). In addition, the CCR5 inhibitor Maraviroc improved the immune response in HNSCC and chordoma (39,68), while a CCR2 inhibitor in combination with bevacizumab reduced tumor size in mouse models of ovarian cancer by inhibiting the MAPK pathway (69). Interestingly, the activation of TAM also fits with an increased presence of IgE. Indeed, anti-tumoral IgEs were shown to increase macrophage inflammatory properties and cytokine production (70). This probably occurs to the FCεRI, which is used to identify TAMs (71). Taken together, the increased expression of immunotherapy targets on myeloid APCs may offer more therapeutic options, and their proximity to a greater CD8^+^ T cell may potentiate the response to immunotherapies in TLS^+^ tumors.

TLS maturation segregated patients into 3 distinct groups, TLS^-^, TLS^imm.^ and TLS^mat.^, and were logically associated with increasing B cell and DC densities and numbers. TLS^mat.^ tumors were associated with improved PFS, compared to TLS^imm.^ and TLS^-^ tumors. These results are in accordance with previous studies showing that TLS maturation is associated with better prognosis (15,41,72) and often helps stratify patients when used as an independent prognostic marker (20) or therapy response marker (60,61). Indeed, TLS presence has refined the molecular classification of patients with endometrial tumors, by sub-stratifying those with MSI into an MSI^+^TLS^+^ patient group with improved prognosis (20). In HNSCC, melanoma and sarcoma, B cell-alone mediated survival rates similar to those seen in TLS^+^ tumors (19,60,61).

In conclusion, our results show that TLS in cervical cancer arise from a microenvironment which is more prone to anti-tumoral response. Indeed, TLS^+^ tumors are enriched in B cells, aDCs, activated macrophages and immune checkpoint positive T cells. Furthermore, activated DCs and macrophages express a greater diversity of costimulatory molecules, are in close proximity of CD8^+^ T cells, which indicates a greater potential for T cell stimulation. Altogether, our results provide a comprehensive characterization of the immune microenvironment in cervical tumors, in terms of phenotype, spatial organization, and immune cell interactions. Furthermore, our results highlight that TLS are associated with well-differentiated cells expressing immunotherapy targets, raising the possibility of TLS as a predictive marker for patient response to specific therapies.

## Declarations

### Ethics approval and consent to participate

Human cervical tumor samples were obtained from patients, who were included in the prospective XAC-03 (NCT02875990) and GC-Bio-IPC (NCT01977274) clinical trials, conducted from 2013 to 2023. The XAC-30 and GC-Bio_IPC trials were approved by the institutional review board (Comité d’Orientation Stratégique (COS), Marseille, France) of the Paoli-Calmettes Institute and the Assistance Publique des Hopitaux de Marseille (AP-HM), respectively. In accordance with the Declaration of Helsinki, written informed consent was obtained from all patients.

### Data Availability

All data presented in the manuscript are original results and are available upon reasonable request to. The publicly available TCGA CESC dataset, used for TLS cell enrichment and signature associated survival, is available on the TCGA data portal https://portal.gdc.cancer.gov/projects/TCGA-CESC.

### Competing interests

D.O. is a cofounder and shareholder of Imcheck Therapeutics, Alderaan Biotechnology, Emergence Therapeutics, and Stealth IO, which did not take part in any part of the manuscript.

RS declares to have received research grants from Astra-Zeneca, consulting fees from GSK and EISAI, and non-financial support from MSD, Astra-Zeneca, GSK, and Novartis, none of which participated to this study.

EL declares to have received consulting fees from GSK and MSD, none of which participated to this study.

Other authors do not have any conflits of interest to declare.

## Acknowledgements

We would like to thank the translational histopathology ICEP platform (CRCM, Marseille, France) for destocking patient FFPE blocks, and cutting the slides. We would like to thank Manon Richaud and Françoise Mallet at the CRCM Cytometry platform (CRCM, Marseille, France) for their insights in panel design and access to cytometers. Editorial assistance for English language improvement was provided by Angloscribe, Calvisson, France.

## Author contributions

L.G. and D.O. designed the project. L.G., J.S., C.D., and N.B., dissociated tumor samples. L.G., M.S.R., N.B., A.B.A., and J.S. performed the phenotyping experiment. L.G., N.B., and O.Q. performed the soluble factor dosages and analysis. M.P. performed the IF staining and acquisitions. E.B performed and interpreted LASSO analysis. M.P. and L.G. performed spatial analyses. L.G. analyzed CyTOF and flow cytometry experiments. X.C., R.S., E.L., and O.Q. enrolled patients in the studies, performed patient follow-up, extracted clinical data and obtained tumor samples. S.G., L.G. and S.F. performed the statistical analyses on cervical tumors. L.G. and A.S.C. performed the statistical analyses and the MFA analysis of clinical data. L.G., M.P. and M.C.D.N. interpreted the data. M.C.D.N. and D.O. supervised the project. L.G. drafted the manuscript and designed the figures. J.A.N aided in manuscript corrections. All authors reviewed the manuscript and provided insights. We declare that all authors contributed to the study and accept accountability of the data herein, and fulfil the authorship criteria.

## Consent for publication

Not applicable

## Additional information

**Correspondence** and requests for materials should be addressed to Laurent Gorvel or Daniel Olive.

---

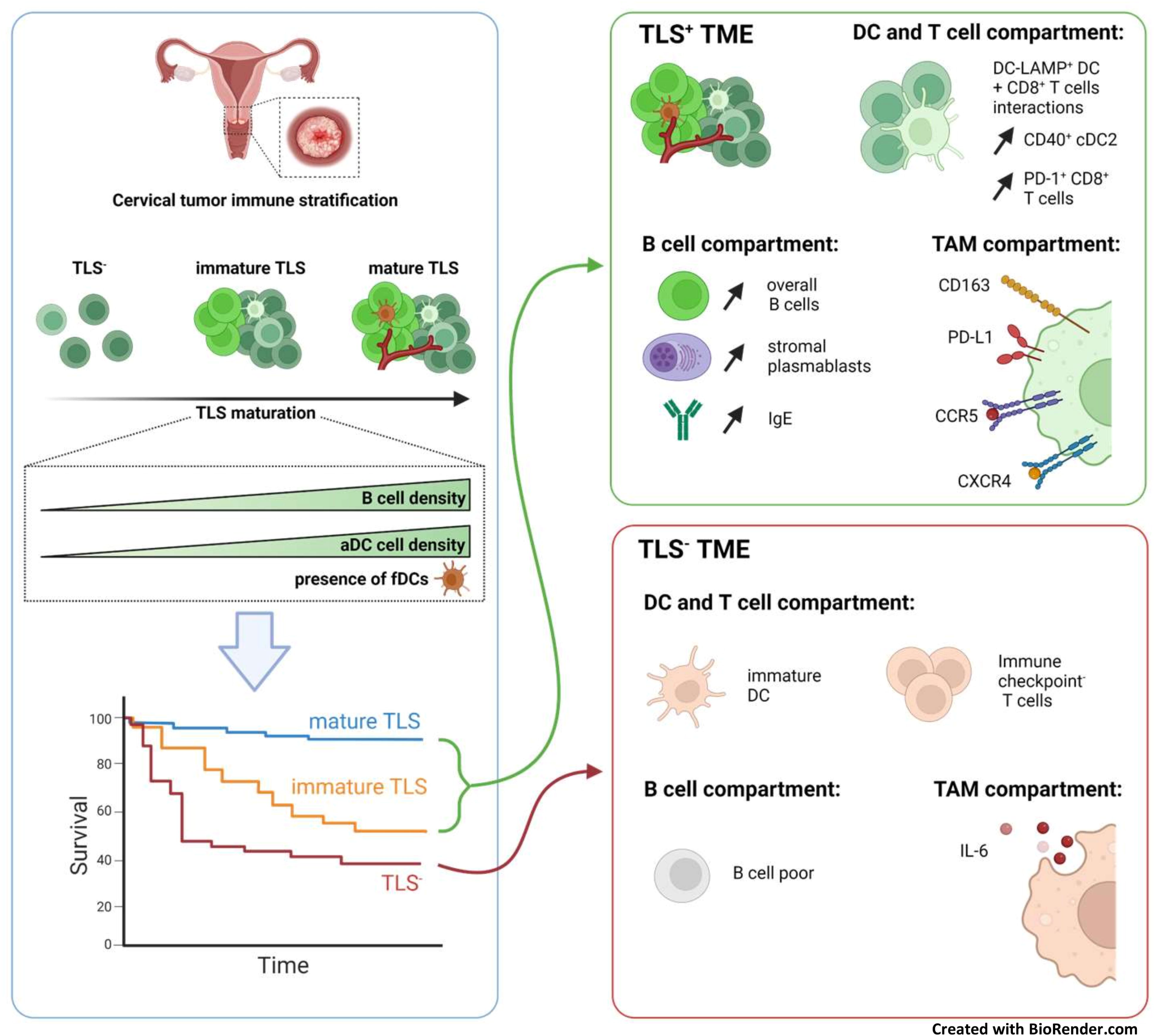

## References

1. Sung H, Ferlay J, Siegel RL, Laversanne M, Soerjomataram I, Jemal A, et al. Global Cancer Statistics 2020: GLOBOCAN Estimates of Incidence and Mortality Worldwide for 36 Cancers in 185 Countries. CA Cancer J Clin 2021;71:209–49

2. Nayar R, Wilbur DC. The Pap test and Bethesda 2014. Cancer Cytopathol 2015;123:271–81

3. Odicino F, Pecorelli S, Zigliani L, Creasman WT. History of the FIGO cancer staging system. Int J Gynaecol Obstet 2008;101:205–10

4. Tong WW, Shepherd K, Garland S, Meagher A, Templeton DJ, Fairley CK, et al. Human papillomavirus 16-specific T-cell responses and spontaneous regression of anal high-grade squamous intraepithelial lesions. The Journal of infectious diseases 2015;211:405–15

5. Koskimaa HM, Paaso AE, Welters MJ, Grenman SE, Syrjanen KJ, van der Burg SH, et al. Human papillomavirus 16 E2-, E6- and E7-specific T-cell responses in children and their mothers who developed incident cervical intraepithelial neoplasia during a 14-year follow-up of the Finnish Family HPV cohort. J Transl Med 2014;12:44

6. Woo YL, van den Hende M, Sterling JC, Coleman N, Crawford RA, Kwappenberg KM, et al. A prospective study on the natural course of low-grade squamous intraepithelial lesions and the presence of HPV16 E2-, E6- and E7-specific T-cell responses. Int J Cancer 2010;126:133-41

7. Cao L, Sun PL, He Y, Yao M, Gao H. Immune stromal features in cervical squamous cell carcinoma are prognostic factors for distant metastasis: A retrospective study. Pathol Res Pract 2020;216:152751

8. Chen XJ, Han LF, Wu XG, Wei WF, Wu LF, Yi HY, et al. Clinical Significance of CD163+ and CD68+ Tumor-associated Macrophages in High-risk HPV-related Cervical Cancer. J Cancer 2017;8:3868–75

9. Guo F, Kong W, Zhao G, Cheng Z, Ai L, Lv J, et al. The correlation between tumor-associated macrophage infiltration and progression in cervical carcinoma. Biosci Rep 2021;41

10. Kawachi A, Yoshida H, Kitano S, Ino Y, Kato T, Hiraoka N. Tumor-associated CD204(+) M2 macrophages are unfavorable prognostic indicators in uterine cervical adenocarcinoma. Cancer Sci 2018;109:863–70

11. Colbert LE, El MB, Lynn EJ, Bronk J, Karpinets TV, Wu X, et al. Expansion of Candidate HPV-Specific T Cells in the Tumor Microenvironment during Chemoradiotherapy Is Prognostic in HPV16(+) Cancers. Cancer Immunol Res 2022;10:259–71

12. Li L, Ma Y, Liu S, Zhang J, Xu XY. Interleukin 10 promotes immune response by increasing the survival of activated CD8(+) T cells in human papillomavirus 16-infected cervical cancer. Tumour Biol 2016

13. Hanna BS, Llao-Cid L, Iskar M, Roessner PM, Klett LC, Wong JKL, et al. Interleukin-10 receptor signaling promotes the maintenance of a PD-1(int) TCF-1(+) CD8(+) T cell population that sustains anti-tumor immunity. Immunity 2021;54:2825–41 e10

14. Schumacher TN, Thommen DS. Tertiary lymphoid structures in cancer. Science 2022;375:eabf9419

15. Fridman WH, Meylan M, Petitprez F, Sun CM, Italiano A, Sautes-Fridman C. B cells and tertiary lymphoid structures as determinants of tumour immune contexture and clinical outcome. Nat Rev Clin Oncol 2022;19:441–57

16. Dieu-Nosjean MC. Tumor-Associated Tertiary Lymphoid Structures: A Cancer Biomarker and a Target for Next-generation Immunotherapy. Adv Exp Med Biol 2021;1329:51–68

17. Sautes-Fridman C, Lawand M, Giraldo NA, Kaplon H, Germain C, Fridman WH, et al. Tertiary Lymphoid Structures in Cancers: Prognostic Value, Regulation, and Manipulation for Therapeutic Intervention. Front Immunol 2016;7:407

18. Lin Z, Huang L, Li S, Gu J, Cui X, Zhou Y. Pan-cancer analysis of genomic properties and clinical outcome associated with tumor tertiary lymphoid structure. Sci Rep 2020;10:21530

19. Ruffin AT, Cillo AR, Tabib T, Liu A, Onkar S, Kunning SR, et al. B cell signatures and tertiary lymphoid structures contribute to outcome in head and neck squamous cell carcinoma. Nat Commun 2021;12:3349

20. Horeweg N, Workel HH, Loiero D, Church DN, Vermij L, Leon-Castillo A, et al. Tertiary lymphoid structures critical for prognosis in endometrial cancer patients. Nat Commun 2022;13:1373

21. Ben Amara A, Rouviere MS, Fattori S, Wlosik J, Gregori E, Boucherit N, et al. High-throughput mass cytometry staining for deep phenotyping of human natural killer cells. STAR Protoc 2022;3:101768

22. van Unen V, Hollt T, Pezzotti N, Li N, Reinders MJT, Eisemann E, et al. Visual analysis of mass cytometry data by hierarchical stochastic neighbour embedding reveals rare cell types. Nat Commun 2017;8:1740

23. Kang J, Choi YJ, Kim IK, Lee HS, Kim H, Baik SH, et al. LASSO-Based Machine Learning Algorithm for Prediction of Lymph Node Metastasis in T1 Colorectal Cancer. Cancer Res Treat 2021;53:773–83

24. Chen B, Khodadoust MS, Liu CL, Newman AM, Alizadeh AA. Profiling Tumor Infiltrating Immune Cells with CIBERSORT. Methods Mol Biol 2018;1711:243–59

25. Guedj K, Khallou-Laschet J, Clement M, Morvan M, Gaston AT, Fornasa G, et al. M1 macrophages act as LTbetaR-independent lymphoid tissue inducer cells during atherosclerosis-related lymphoid neogenesis. Cardiovasc Res 2014;101:434–43

26. Kang W, Feng Z, Luo J, He Z, Liu J, Wu J, et al. Tertiary Lymphoid Structures in Cancer: The Double-Edged Sword Role in Antitumor Immunity and Potential Therapeutic Induction Strategies. Front Immunol 2021;12:689270

27. Goc J, Germain C, Vo-Bourgais TK, Lupo A, Klein C, Knockaert S, et al. Dendritic cells in tumor-associated tertiary lymphoid structures signal a Th1 cytotoxic immune contexture and license the positive prognostic value of infiltrating CD8+ T cells. Cancer Res 2014;74:705–15

28. Coppola D, Nebozhyn M, Khalil F, Dai H, Yeatman T, Loboda A, et al. Unique ectopic lymph node-like structures present in human primary colorectal carcinoma are identified by immune gene array profiling. Am J Pathol 2011;179:37–45

29. Fernandez-Avila L, Castro-Amaya AM, Molina-Pineda A, Hernandez-Gutierrez R, Jave-Suarez LF, Aguilar-Lemarroy A. The Value of CXCL1, CXCL2, CXCL3, and CXCL8 as Potential Prognosis Markers in Cervical Cancer: Evidence of E6/E7 from HPV16 and 18 in Chemokines Regulation. Biomedicines 2023;11

30. de Chaisemartin L, Goc J, Damotte D, Validire P, Magdeleinat P, Alifano M, et al. Characterization of chemokines and adhesion molecules associated with T cell presence in tertiary lymphoid structures in human lung cancer. Cancer Res 2011;71:6391–9

31. Liu Y, Cruikshank WW, O’Loughlin T, O’Reilly P, Center DM, Kornfeld H. Identification of a CD4 domain required for interleukin-16 binding and lymphocyte activation. J Biol Chem 1999;274:23387-95

32. Lynch EA, Heijens CA, Horst NF, Center DM, Cruikshank WW. Cutting edge: IL-16/CD4 preferentially induces Th1 cell migration: requirement of CCR5. Journal of immunology 2003;171:4965–8

33. Kuo P, Tuong ZK, Teoh SM, Frazer IH, Mattarollo SR, Leggatt GR. HPV16E7-Induced Hyperplasia Promotes CXCL9/10 Expression and Induces CXCR3(+) T-Cell Migration to Skin. J Invest Dermatol 2018;138:1348–59

34. Liang YK, Deng ZK, Chen MT, Qiu SQ, Xiao YS, Qi YZ, et al. CXCL9 Is a Potential Biomarker of Immune Infiltration Associated With Favorable Prognosis in ER-Negative Breast Cancer. Front Oncol 2021;11:710286

35. Li Z, Liu J, Li L, Shao S, Wu J, Bian L, et al. Epithelial mesenchymal transition induced by the CXCL9/CXCR3 axis through AKT activation promotes invasion and metastasis in tongue squamous cell carcinoma. Oncol Rep 2018;39:1356–68

36. Li M, He L, Zhu J, Zhang P, Liang S. Targeting tumor-associated macrophages for cancer treatment. Cell Biosci 2022;12:85

37. Zhang XF, Zhang XL, Wang YJ, Fang Y, Li ML, Liu XY, et al. The regulatory network of the chemokine CCL5 in colorectal cancer. Ann Med 2023;55:2205168

38. Aldinucci D, Borghese C, Casagrande N. The CCL5/CCR5 Axis in Cancer Progression. Cancers (Basel) 2020;12

39. Xu J, Shi Q, Lou J, Wang B, Wang W, Niu J, et al. Chordoma recruits and polarizes tumor-associated macrophages via secreting CCL5 to promote malignant progression. J Immunother Cancer 2023;11

40. Cambien B, Richard-Fiardo P, Karimdjee BF, Martini V, Ferrua B, Pitard B, et al. CCL5 neutralization restricts cancer growth and potentiates the targeting of PDGFRbeta in colorectal carcinoma. PloS one 2011;6:e28842

41. Dieu-Nosjean MC, Antoine M, Danel C, Heudes D, Wislez M, Poulot V, et al. Long-term survival for patients with non-small-cell lung cancer with intratumoral lymphoid structures. J Clin Oncol 2008;26:4410–7

42. Karapetyan L, AbuShukair HM, Li A, Knight A, Al Bzour AN, MacFawn IP, et al. Expression of lymphoid structure-associated cytokine/chemokine gene transcripts in tumor and protein in serum are prognostic of melanoma patient outcomes. Front Immunol 2023;14:1171978

43. Li C, Hua K. Dissecting the Single-Cell Transcriptome Network of Immune Environment Underlying Cervical Premalignant Lesion, Cervical Cancer and Metastatic Lymph Nodes. Front Immunol 2022;13:897366

44. Chauhan J, Grandits M, Palhares L, Mele S, Nakamura M, Lopez-Abente J, et al. Anti-cancer pro-inflammatory effects of an IgE antibody targeting the melanoma-associated antigen chondroitin sulfate proteoglycan 4. Nat Commun 2023;14:2192

45. Yao X, Chen W, Zhao C, Wei L, Hu Y, Li M, et al. Naturally acquired HPV antibodies against subsequent homotypic infection: A large-scale prospective cohort study. Lancet Reg Health West Pac 2021;13:100196

46. Beachler DC, Jenkins G, Safaeian M, Kreimer AR, Wentzensen N. Natural Acquired Immunity Against Subsequent Genital Human Papillomavirus Infection: A Systematic Review and Meta-analysis. The Journal of infectious diseases 2016;213:1444–54

47. Safaeian M, Castellsague X, Hildesheim A, Wacholder S, Schiffman MH, Bozonnat MC, et al. Risk of HPV-16/18 Infections and Associated Cervical Abnormalities in Women Seropositive for Naturally Acquired Antibodies: Pooled Analysis Based on Control Arms of Two Large Clinical Trials. The Journal of infectious diseases 2018;218:84–94

48. Gorvel L, Olive D. Tumor associated macrophage in HPV(+) tumors: Between immunosuppression and inflammation. Semin Immunol 2022;65:101671

49. Pittet MJ, Michielin O, Migliorini D. Clinical relevance of tumour-associated macrophages. Nat Rev Clin Oncol 2022;19:402–21

50. Park MD, Silvin A, Ginhoux F, Merad M. Macrophages in health and disease. Cell 2022;185:4259–79

51. van Vlerken-Ysla L, Tyurina YY, Kagan VE, Gabrilovich DI. Functional states of myeloid cells in cancer. Cancer Cell 2023;41:490–504

52. Del Prete A, Salvi V, Soriani A, Laffranchi M, Sozio F, Bosisio D, et al. Dendritic cell subsets in cancer immunity and tumor antigen sensing. Cell Mol Immunol 2023;20:432–47

53. Truxova I, Kasikova L, Hensler M, Skapa P, Laco J, Pecen L, et al. Mature dendritic cells correlate with favorable immune infiltrate and improved prognosis in ovarian carcinoma patients. J Immunother Cancer 2018;6:139

54. Minohara K, Imai M, Matoba T, Wing JB, Shime H, Odanaka M, et al. Mature dendritic cells enriched in regulatory molecules may control regulatory T cells and the prognosis of head and neck cancer. Cancer Sci 2023;114:1256–69

55. Duraiswamy J, Turrini R, Minasyan A, Barras D, Crespo I, Grimm AJ, et al. Myeloid antigen-presenting cell niches sustain antitumor T cells and license PD-1 blockade via CD28 costimulation. Cancer Cell 2021;39:1623–42 e20

56. Haque A, Moriyama M, Kubota K, Ishiguro N, Sakamoto M, Chinju A, et al. CD206(+) tumor-associated macrophages promote proliferation and invasion in oral squamous cell carcinoma via EGF production. Sci Rep 2019;9:14611

57. Yagi T, Baba Y, Okadome K, Kiyozumi Y, Hiyoshi Y, Ishimoto T, et al. Tumour-associated macrophages are associated with poor prognosis and programmed death ligand 1 expression in oesophageal cancer. Eur J Cancer 2019;111:38–49

58. Li Z, Maeda D, Yoshida M, Umakoshi M, Nanjo H, Shiraishi K, et al. The intratumoral distribution influences the prognostic impact of CD68- and CD204-positive macrophages in non-small cell lung cancer. Lung Cancer 2018;123:127–35

59. Schweer D, McAtee A, Neupane K, Richards C, Ueland F, Kolesar J. Tumor-Associated Macrophages and Ovarian Cancer: Implications for Therapy. Cancers (Basel) 2022;14

60. Petitprez F, de Reynies A, Keung EZ, Chen TW, Sun CM, Calderaro J, et al. B cells are associated with survival and immunotherapy response in sarcoma. Nature 2020;577:556–60

61. Helmink BA, Reddy SM, Gao J, Zhang S, Basar R, Thakur R, et al. B cells and tertiary lymphoid structures promote immunotherapy response. Nature 2020;577:549–55

62. Wang L, Guo W, Guo Z, Yu J, Tan J, Simons DL, et al. PD-L1-expressing tumor-associated macrophages are immunostimulatory and associate with good clinical outcome in human breast cancer. Cell Rep Med 2024;5:101420

63. Zhang Y, Wang N, Ding M, Yang Y, Wang Z, Huang L, et al. CD40 Accelerates the Antigen-Specific Stem-Like Memory CD8(+) T Cells Formation and Human Papilloma Virus (HPV)-Positive Tumor Eradication. Front Immunol 2020;11:1012

64. Ceglia V, Zurawski S, Montes M, Kroll M, Bouteau A, Wang Z, et al. Anti-CD40 Antibody Fused to CD40 Ligand Is a Superagonist Platform for Adjuvant Intrinsic DC-Targeting Vaccines. Front Immunol 2021;12:786144

65. Vitale LA, Thomas LJ, He LZ, O’Neill T, Widger J, Crocker A, et al. Development of CDX-1140, an agonist CD40 antibody for cancer immunotherapy. Cancer Immunol Immunother 2019;68:233–45

66. Labiano S, Roh V, Godfroid C, Hiou-Feige A, Romero J, Sum E, et al. CD40 Agonist Targeted to Fibroblast Activation Protein alpha Synergizes with Radiotherapy in Murine HPV-Positive Head and Neck Tumors. Clin Cancer Res 2021;27:4054–65

67. Nywening TM, Wang-Gillam A, Sanford DE, Belt BA, Panni RZ, Cusworth BM, et al. Targeting tumour-associated macrophages with CCR2 inhibition in combination with FOLFIRINOX in patients with borderline resectable and locally advanced pancreatic cancer: a single-centre, open-label, dose-finding, non-randomised, phase 1b trial. Lancet Oncol 2016;17:651–62

68. Dunbar KJ, Karakasheva TA, Tang Q, Efe G, Lin EW, Harris M, et al. Tumor-Derived CCL5 Recruits Cancer-Associated Fibroblasts and Promotes Tumor Cell Proliferation in Esophageal Squamous Cell Carcinoma. Mol Cancer Res 2023;21:741–52

69. Zhai T, Mitamura T, Wang L, Kubota SI, Murakami M, Tanaka S, et al. Combination therapy with bevacizumab and a CCR2 inhibitor for human ovarian cancer: An in vivo validation study. Cancer Med 2023;12:9697–708

70. Pellizzari G, Hoskin C, Crescioli S, Mele S, Gotovina J, Chiaruttini G, et al. IgE re-programs alternatively-activated human macrophages towards pro-inflammatory anti-tumoural states. EBioMedicine 2019;43:67–81

71. Dong K, Chen W, Pan X, Wang H, Sun Y, Qian C, et al. FCER1G positively relates to macrophage infiltration in clear cell renal cell carcinoma and contributes to unfavorable prognosis by regulating tumor immunity. BMC Cancer 2022;22:140

72. Sautes-Fridman C, Verneau J, Sun CM, Moreira M, Chen TW, Meylan M, et al. Tertiary Lymphoid Structures and B cells: Clinical impact and therapeutic modulation in cancer. Semin Immunol 2020;48:101406

