## Supplementary material for "Tertiary lymphoid structures are associated with enhanced macrophage activation, immune checkpoint expression and predict outcome in cervical cancer": Supp Figures

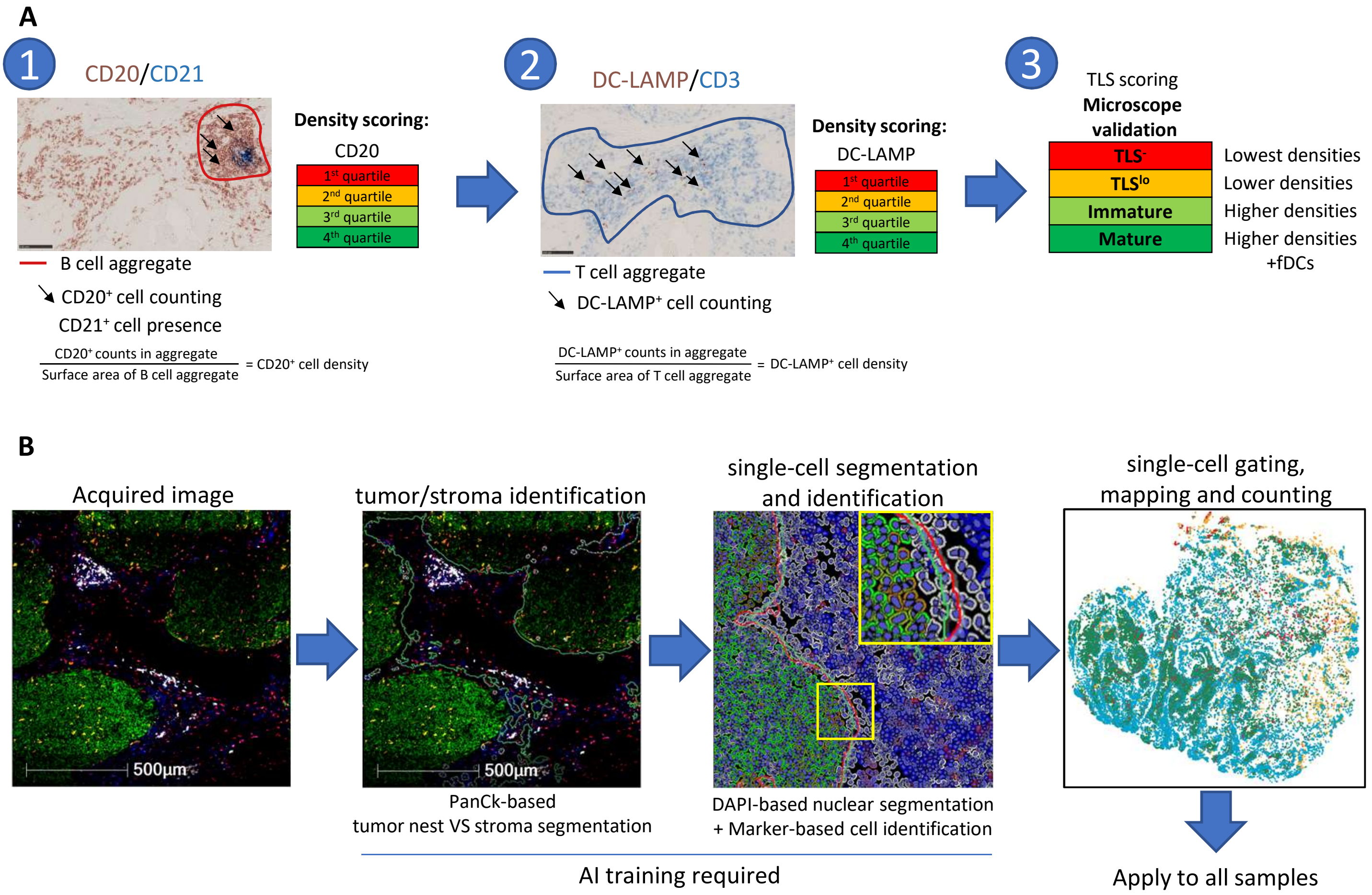


**Supp. Fig. 1: TLS maturation classification and cell segmentation by multiplex IHC. A.** Immunohistochemistry (IHC) stained cervical tumor samples (n=34) were screened for $CD3^+$ T cell aggregates (blue circling). B cells ($CD20^+$) inside B cell aggregates and activated DCs (DC-$LAMP^+$) located in T cell ($CD3^+$) aggregates were counted. Their density was assessed by dividing B cell or activated DC count by the surface of the aggregate (in $µm^2$). The obtained B cell and activated DC densities (in count/$µm^2$) were then scored by quartile, and combined to assess TLS presence. A higher score for B cell and activated DC density resulted in higher TLS score (quartile) and more formed TLS while a lower TLS score resulted in absence of TLS or unorganized aggregates. TLS presence and maturity were then confirmed by microscopy and CD21 staining marking Follicular DCs (fDCs). **B**, Multiplex IHC images were analyzed to identify tumor nests ($PanCk^+$ regions of the sample, green) or stroma (PanCk- regions of the sample). This was assessed by training an AI-based algorithm to recognize such regions across samples. Following identification of tissue regions, single cell annotation was performed. AI-based algorithm was trained to recognize cell nuclei based on DAPI staining, and to annotate cells based on marker expression, for example CD8+ T cells (PanCk-$CD3^+CD8^+$, in red). This identification was applied across samples and allowed cell subset recognition, mapping on the entire sample and counting.

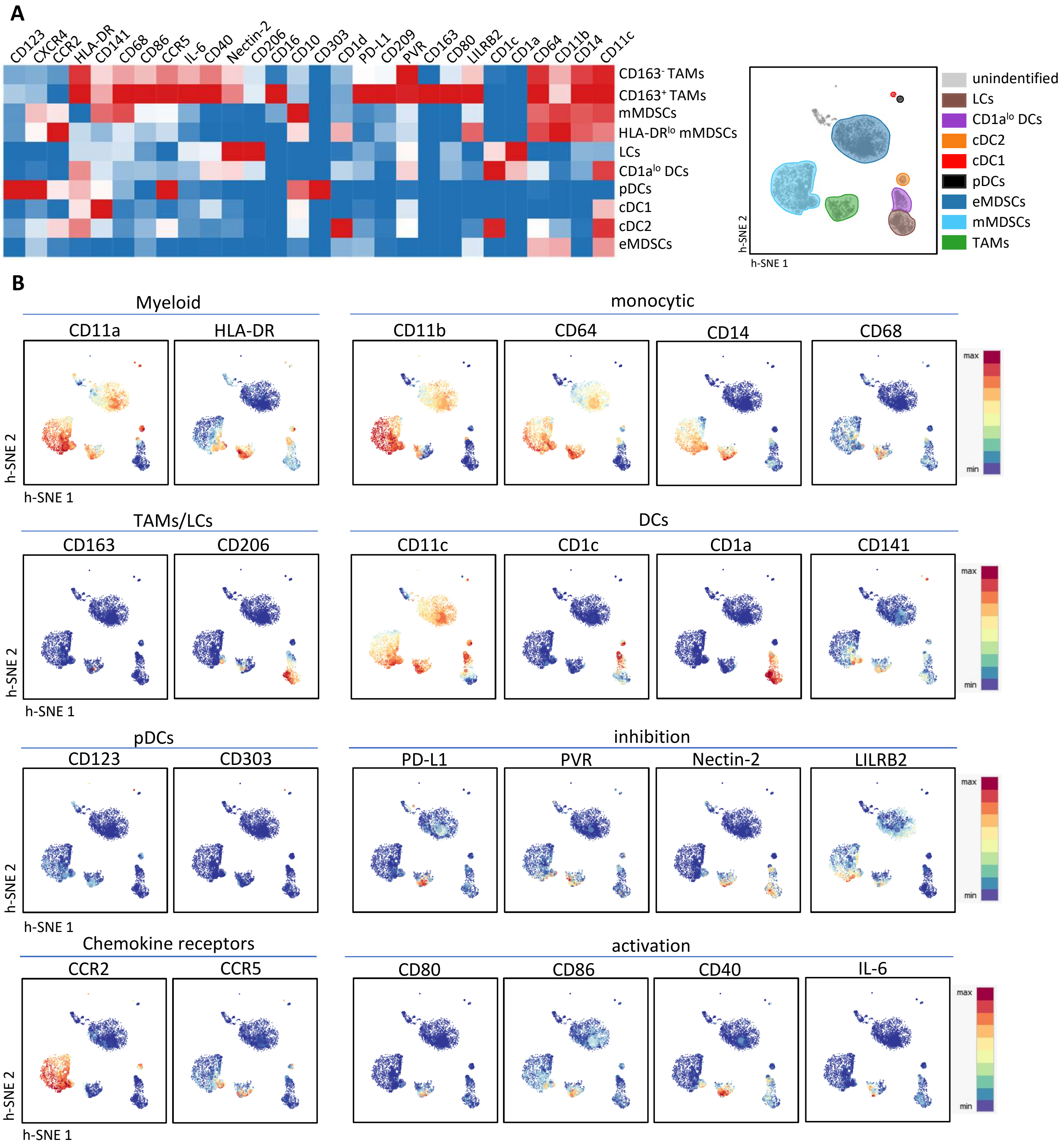


**Supp. Figure 2: Myeloid cell cluster annotation within cervical tumors. A**, Heatmap represents the expression of markers against different myeloid cell subsets which were reported on the h-SNE map. **B**, Each marker expression was projected on the h-SNE map.

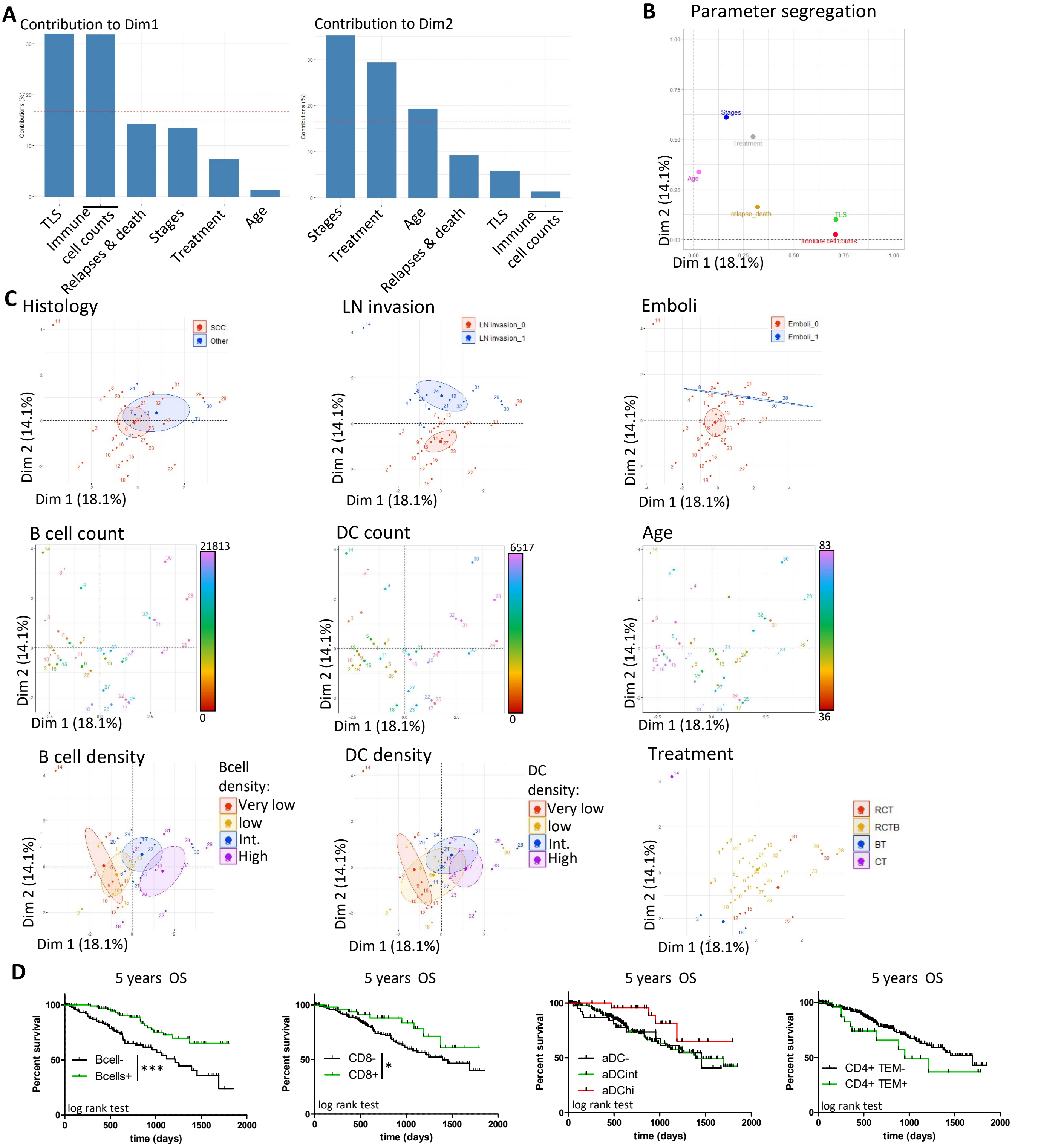


**Supplementary Figure 3: MFA individual parameters and immune cell enrichment-related survival.** Multiple factor analysis (MFA) was run on 34 samples according to TLS maturation status, B cell density, DC density, B cell count, DC count, Emboli presence, LN invasion, Histology, Age, Treatment, relapses and death, and FIGO status. For to allow figure readability certain parameters were grouped. TLS: TLS maturation + B cell density + DC density. Immune cell counts: B cell counts + DC counts. Stages: FIGO stages + Histology + LN invasion + Emboli. **A.** Contribution of each parameter and grouped parameters to dimensions. **B.** Segregation plots of the different parameters **C.** MFA plots patient repartition for Histology, LN invasion, Emboli, B cell counts, DC counts, Age, B cell density, DC density and Treatment. Circles show 95% confidence index. **D**, Immune cell enrichment algorithm Cibersortx was used to investigate TIL infiltrate in TCGA CESC cohort (309 patients). Kaplan-Meier curves were generated for B cells (B cell$^+$ in green, Bcell$^-$ in black), CD8$^+$ Tcells (CD8$^+$ in green, CD8$^-$ in black), activated DCs (aDC$^{hi}$ in red, aDC$^{int}$ in green, aDC$^-$ in black) and CD4+ TEM (CD4$^+$TEM$^+$ in green, CD4$^+$TEM$^-$ in black. *,*p*<0.05 and ***, *p*<0.005.

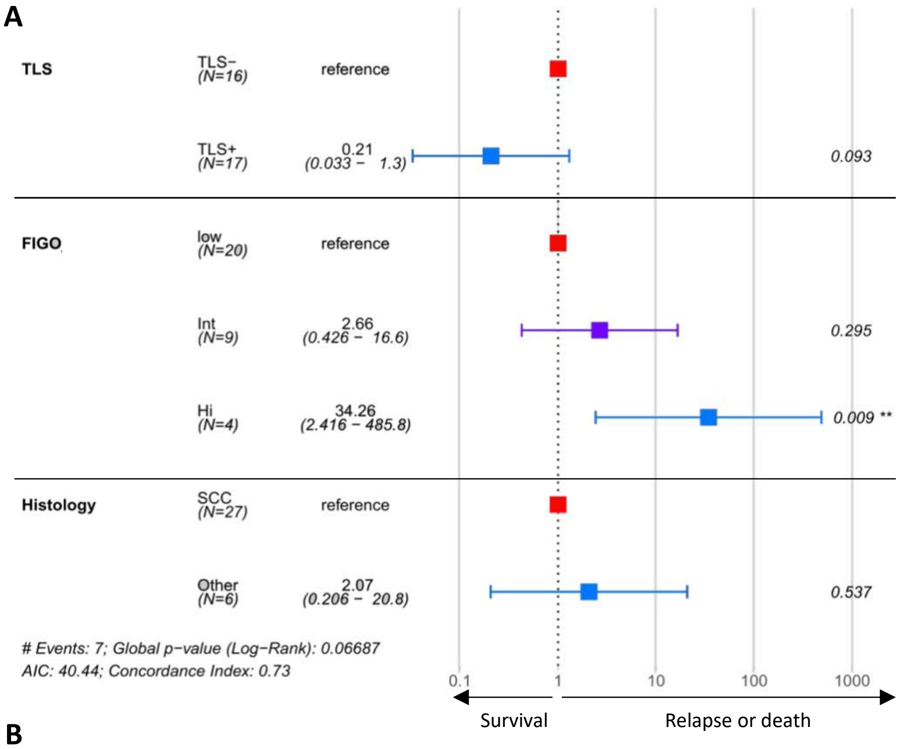


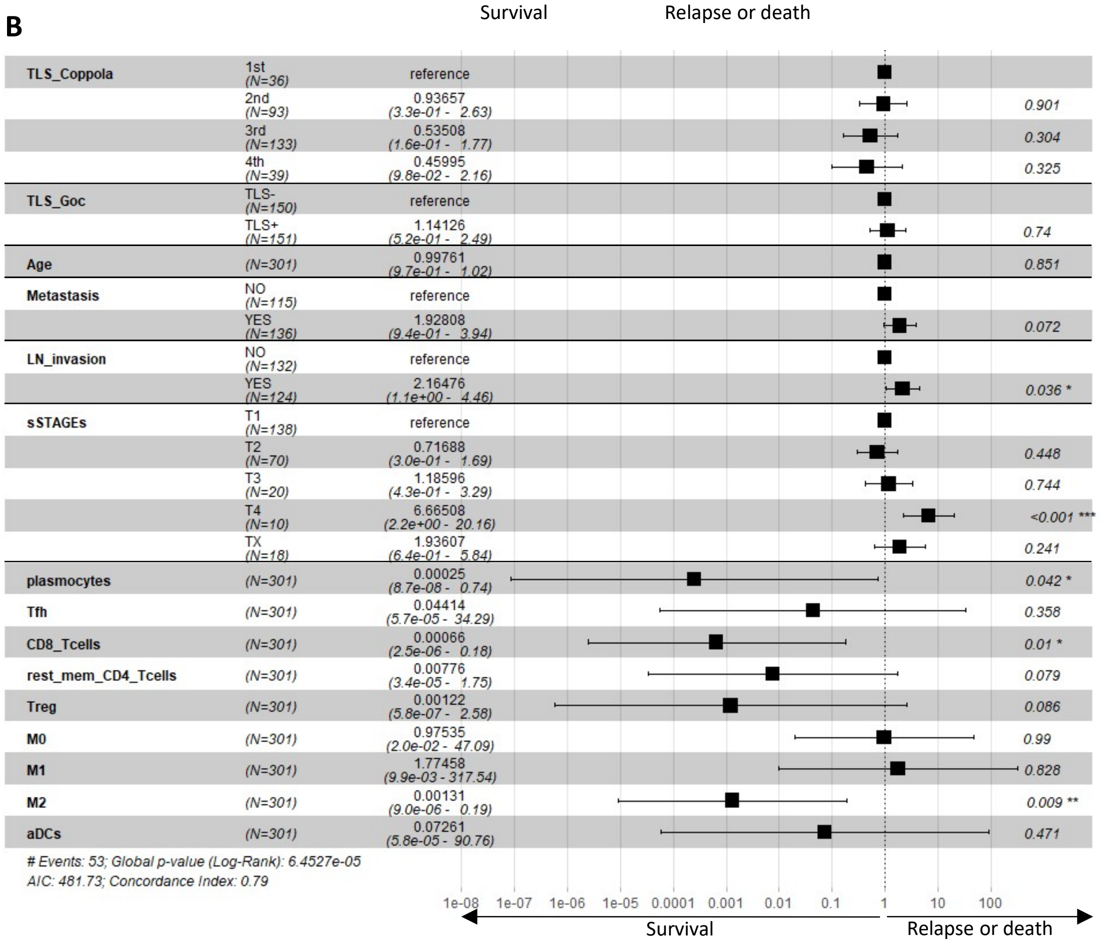


Supplementary Figure 4

**Supplementary Figure 4: Multivariate analysis of cervical cancer patient clinical and TLS parameters**. Multivariate analysis was conducted on n=34 patients. Input parameters were the presence or absence of TLS ($TLS^+$ and $TLS^-$), low FIGO stages (FIGO I and FIGO II, local-regional tumors), Intermediate FIGO stages (FIGO III, LN invasion), High FIGO stages (FIGO IV, metastatic, systemic invaison) and Histology (SCC or other tumor types). **B**. Multivariate analysis was conducted on the CESC TCGA cohort. TLS signatures are obained from *Coppola et al.* and *Goc et al.*. Stages ranked from T1 to T4 and TX, individual population scores were obtained from CibersortX enrichment. For both analyses, cox regression was performed and Log-rank test calculated the global and individual *p*-values. **p*-value<0.05
