## Supplementary material for "Tertiary lymphoid structures are associated with enhanced macrophage activation, immune checkpoint expression and predict outcome in cervical cancer": Supp File 1

***#AFM - Analyse Factorielle Multiple avec R: L'Essentiel***

install.packages(c("FactoMineR", "factoextra"))

library("FactoMineR")

library("factoextra")

setwd("D:/TLS paper ongoing/MFA paper version/AFM paper version")

AFM <- Data3

class("TLS")

***# MFA***

#MFA (base, group, type = rep("s",length(group)), ind.sup = NULL,

\# name.group = NULL, num.group.sup = NULL, graph = TRUE)

*#with all parameters*

res.mfa <- MFA(AFM[,-1],

group = c( 1, 1, 1, 1, 1, 1, 1, 1, 1, 1, 1, 1),

type = c( "c", "c", "n", "n", "n","n", "n", "n","n","n", "c","c"),

name.group = c("DC.number", "Bcell.number", "DC.density","Bcell.Density", "TLS",

"FDC","Histology", "LN invasion", "Emboli", "FIGO stages", "Age", "Age_diagnosis"),

num.group.sup = NULL, graph = TRUE)

print(res.mfa)

***# Groups***

res.mfag <- MFA(AFM[,-1],

group = c(2,2, 1, 1, 3, 1,2),

type = c("c","n", "n", "n", "n", "n","c"),

name.group = c("Immune cell counts","Immune cell densities", "TLS", "FDC", "Pathology", "Stages","Age"),

num.group.sup = NULL,

graph = TRUE)

print(res.mfag)

***# Variances***

library("factoextra")

eig.val <- get_eigenvalue(res.mfag)

head(eig.val)

fviz_screeplot(res.mfa)

***# Variables graph***

group <- get_mfa_var(res.mfa, "group")

group

***# Group coordinates***

head(group$coord)

*## Cos2: group representation quality*

head(group$cos2)

***# Dimension contribution***

head(group$contrib)

fviz_mfa_var(res.mfa, "group")

*## First dimension contribution*

fviz_contrib (res.mfa, "group", axes = 1)

*## Second dimension contribution*

fviz_contrib (res.mfa, "group", axes = 2)

*#quantitative variable*

quanti.var <- get_mfa_var(res.mfa, "quanti.var")

quanti.var

***#Patient graphs***

*## Multiple correspondence analysis for qualitative variables*

ind <- get_mfa_ind(res.mfa)

ind

*##2 parameters*

fviz_mfa_ind(res.mfa,

habillage = "FDC", # color by groups

palette = c("#F03209", "#0644E8"),

addEllipses = TRUE, ellipse.type = "confidence",

repel = TRUE )

*##3 parameters*

fviz_mfa_ind(res.mfa,

habillage = "Histology", # color by groups

palette = c("#F03209", "#E8AA06", "#0644E8"),

addEllipses = TRUE, ellipse.type = "confidence",

repel = TRUE )

*##4 parameters*

fviz_mfa_ind(res.mfa,

habillage = "DC.density", # color by groups

palette = c("#F03209", "#E8AA06", "#0644E8", "#AF16F2"),

addEllipses = TRUE, ellipse.type = "confidence",

repel = TRUE )

***#Parameter visualization***

fviz_ellipses(res.mfa, c("DC.number", "Bcell.number", "Age", "Age_diagnosis"), repel = T)

fviz_ellipses(res.mfa, c("FIGO stages"), repel = T)
