## Supplementary material for "Tertiary lymphoid structures are associated with enhanced macrophage activation, immune checkpoint expression and predict outcome in cervical cancer": Supp Table 1

**Suppl. Table 1. Reagents and resources for immunohistochemistry and immunofluorescence assays**

| **Product** | **Reference** | **Company** | **Final concentration/dilution** |
| --- | --- | --- | --- |
| AEC | SK-4200 | Vector Laboratories | Ready-to-use |
| Biotin-XX Tyramide | B40951 | ThermoFisher Scientific | 1:125 final dilution in 0.03% H_2_0_2_-TBS |
| CF430 Tyramide | 96053 | Interchim | 40 µg/mL in 0.03% H_2_0_2_-TBS |
| CF514 Tyramide | 92199 | Interchim | 10 µg/mL in 0.03% H_2_0_2_-TBS |
| DAPI | 62248 | ThermoFisher Scientific | 1 µg/mL |
| Fluorescence Mounting Medium | S302380-2 | Dako | Ready-to-use |
| Glycergel | C0563 | Dako | Ready-to-use |
| Hydrogen Peroxide H_2_0_2_ (30%) | BP2633-500 | Fisher Bioreagents | 3% in deionized H_2_0 |
| Protein Block, Serum-Free Solution | X090930-2 | Dako | Ready-to-use |
| REAL Antibody Diluent | S202230-2 | Dako | Ready-to-use |
| SAP | SK-5300 | Vector Laboratories | Ready-to-use |
| Streptavidine-AF647 | 405237 | BioLegend | 1 µg/mL |
| Streptavidin-HRP | P0397 | Dako | 2.5 µg/mL |
| Target Retrieval solution high pH | S236784-2 | Dako | 1:10 final dilution in distilled water |
| Target Retrieval solution low pH | S169984-2 | Dako | 1:10 final dilution in distilled water |
| TBS 10X | ET220-B | Euromedex | 1:10 final dilution in in deionized H_2_0 |
| Tween 20 | 1003187960 | Sigma | 0.04% in TBS |
| Tyramide Alexa Fluor™ 488 | B40953 | ThermoFisher Scientific | 1:125 final dilution in 0.03% H_2_0_2_-TBS |
| Tyramide Alexa Fluor™ 555 | B40955 | ThermoFisher Scientific | 1:125 final dilution in 0.03% H_2_0_2_-TBS |
| Tyramide Alexa Fluor™ 594 | B40957 | ThermoFisher Scientific | 1:125 final dilution in 0.03% H_2_0_2_-TBS |
| Tyramide Alexa Fluor™ 647 | B40958 | ThermoFisher Scientific | 1:125 final dilution in 0.03% H_2_0_2_-TBS |
