## Supplementary material for "Tertiary lymphoid structures are associated with enhanced macrophage activation, immune checkpoint expression and predict outcome in cervical cancer": Supp Table 2

**Suppl. Table 2. Primary antibodies used in the immunohistochemistry and immunofluorescence assays**

| Target | Clone | Host/Isotype | Reference | Company | Final concentration/dilution | pH retrieval | Incubation |
| --- | --- | --- | --- | --- | --- | --- | --- |
| CD1a | 010 | Mouse IgG1 | IR069 | Dako | ready-to-use | 6 | ON / 4°C |
| CD3 | polyclonal | Rabbit | A0452 | Dako | 8 µg/mL | 6 | 1h / RT |
| CD8 | C8/144B | Mouse IgG1 | M7103 | Dako | 5.23 µg/mL | 9 | ON / 4°C |
| CD20 | L26 | Mouse IgG2a | M0755 | Dako | 0.504 µg/mL | 6 | 1h / RT |
| CD21 | 1F8 | Mouse IgG1 | M0784 | Dako | 6.6 µg/mL | 6 | ON / 4°C |
| CD68 | KP1 | Mouse IgG1 | IS609 | Dako | 3.7 µg/mL | 6 | 1h / RT |
| CD138 | MI15 | Mouse IgG1 | GA642 | Dako | ready-to-use | 9 | ON / 4°C |
| CD163 | 10D6 | Mouse IgG1 | MA5-11458 | ThermoFisher Scientific | 1:25 final dilution | 6 | 1h / RT |
| DC-Lamp | 1010E1.01 | Rat IgG2a | DDX0191P-100 | Eurobio Scientific | 6.25 µg/mL | 6 | 1h / RT |
| Pan-Cytokeratins | AE1-AE3 | Mouse IgG1 | M3515 | Dako | 3.33 µg/mL | 6 | 1h / RT |
| PNAd | MECA-79 | Rat IgM | 553863 | BD Pharmingen | 10 µg/mL | 6 | 1h / RT |
