## Supplementary material for "Tertiary lymphoid structures are associated with enhanced macrophage activation, immune checkpoint expression and predict outcome in cervical cancer": Supp Table 3

**Suppl. Table 3. Secondary antibodies used in the immunohistochemistry and immunofluorescence assays**

| Secondary antibody | Reference | Company | Final concentration/dilution |
| --- | --- | --- | --- |
| F(ab)'_2_ donkey anti-rabbit IgG - AP | 711-056-152 | Jackson ImmunoResearch | 5 µg/mL |
| F(ab)'_2_ donkey anti-rabbit IgG - HRP | 711-036-152 | Jackson ImmunoResearch | 1.6 µg/mL |
| F(ab)'_2_ donkey anti-rat IgG - Biotin | 712-066-153 | Jackson ImmunoResearch | 2.4 µg/mL |
| F(ab)'_2_ goat anti-rat IgM – AF647 | 112-606-075 | Jackson ImmunoResearch | 15 µg/mL |
| IgG goat anti-mouse IgG1 - AP | 115-055-205 | Jackson ImmunoResearch | 5 µg/mL |
| IgG goat anti-mouse IgG1 - HRP | 115-035-205 | Jackson ImmunoResearch | 1.6 µg/mL |
| IgG goat anti-mouse IgG2a – Biotin | 115-065-206 | Jackson ImmunoResearch | 13 µg/mL |
| IgG goat anti-mouse IgG2a – BV480 | 115-685-206 | Jackson ImmunoResearch | 4 µg/mL |
| IgG goat anti-mouse IgG2a - HRP | 115-035-206 | Jackson ImmunoResearch | 1.6 µg/mL |
| IgG goat anti-rat IgG - HRP | MP-7404 | Vector Laboratories | Ready to use |
