## Supplementary material for "Tertiary lymphoid structures are associated with enhanced macrophage activation, immune checkpoint expression and predict outcome in cervical cancer": Supp Table 4

| **Suppl. Table 4: Mass cytometry myeloid cell phenotyping panel** | | |
| --- | --- | --- |
| Antibody target | Weight | Metal isotope |
| CD11b | 209 | Bi |
| CD80 | 161 | Dy |
| CD209 | 162 | Dy |
| CD33 | 163 | Dy |
| CD15 | 164 | Dy |
| CD1a | 167 | Er |
| CD206 | 168 | Er |
| CD3 | 170 | Er |
| CD24 | 151 | Eu |
| CD192 | 153 | Eu |
| CD1c | 155 | Gd |
| CD184 | 156 | Gd |
| CD10 | 158 | Gd |
| CD40 | 165 | Ho |
| CD163 | 115 | In |
| PD-L1 | 175 | Lu |
| CD11a | 142 | Nd |
| CD123 | 143 | Nd |
| CD195 | 144 | Nd |
| CD163 | 145 | Nd |
| CD64 | 146 | Nd |
| CD141 | 148 | Nd |
| CD86 | 150 | Nd |
| CD303 | 147 | Sm |
| CD141 | 149 | Sm |
| CD11c | 159 | Tb |
| CD1d | 169 | Tm |
| CD45 | 89 | Y |
| LILRB2 (ILT4) | 171 | Yb |
| HLA-DR | 173 | Yb |
| PVR | 174 | Yb |
| NECTIN-2 | 176 | Yb |
| IL-6 | 154 | Sm |
| CD68 | 141 | Pr |
| Cis-Platin | 195 | Pt |
