## Supplementary material for "Tertiary lymphoid structures are associated with enhanced macrophage activation, immune checkpoint expression and predict outcome in cervical cancer": Supp Table 5

| **Suppl. Table 5: Mass cytometry lymphoid cell phenotyping panel** | | |
| --- | --- | --- |
| **Antibody target** | **Weight** | **Metal isotope** |
| CD45RA | 143 | Nd |
| HVEM | 144 | Nd |
| CD8 | 146 | Nd |
| CD25 | 149 | Sm |
| OX40 | 150 | Nd |
| ICOS | 151 | Eu |
| TIM3 | 153 | Eu |
| CD3 | 115 | In |
| TIGIT | 154 | Sm |
| PD-1 | 155 | Gd |
| PD-L1 | 156 | Gd |
| 4-1BB | 158 | Gd |
| CCR7 | 159 | Tb |
| CD28 | 160 | Gd |
| CTLA-4 | 161 | Dy |
| BTLA | 163 | Dy |
| CD95 | 164 | Dy |
| CD127 | 165 | Ho |
| CD44 | 166 | Di |
| CD27 | 167 | Er |
| CD69 | 168 | Er |
| TCR-vd2 | 169 | Tm |
| CD33 | 170 | Er |
| DNAM-1 | 171 | Yb |
| CD57 | 172 | Yb |
| CD4 | 174 | Yb |
| LAG3 | 175 | Lu |
| CD45 | 89 | Y |
| CD56 | 176 | Yb |
| CD16 | 209 | Bi |
| Cis-Platin | 195 | Pt |
