## Supplementary figures and images for "Tertiary lymphoid structures are associated with enhanced macrophage activation, immune checkpoint expression and predict outcome in cervical cancer"

### Supp Table 9

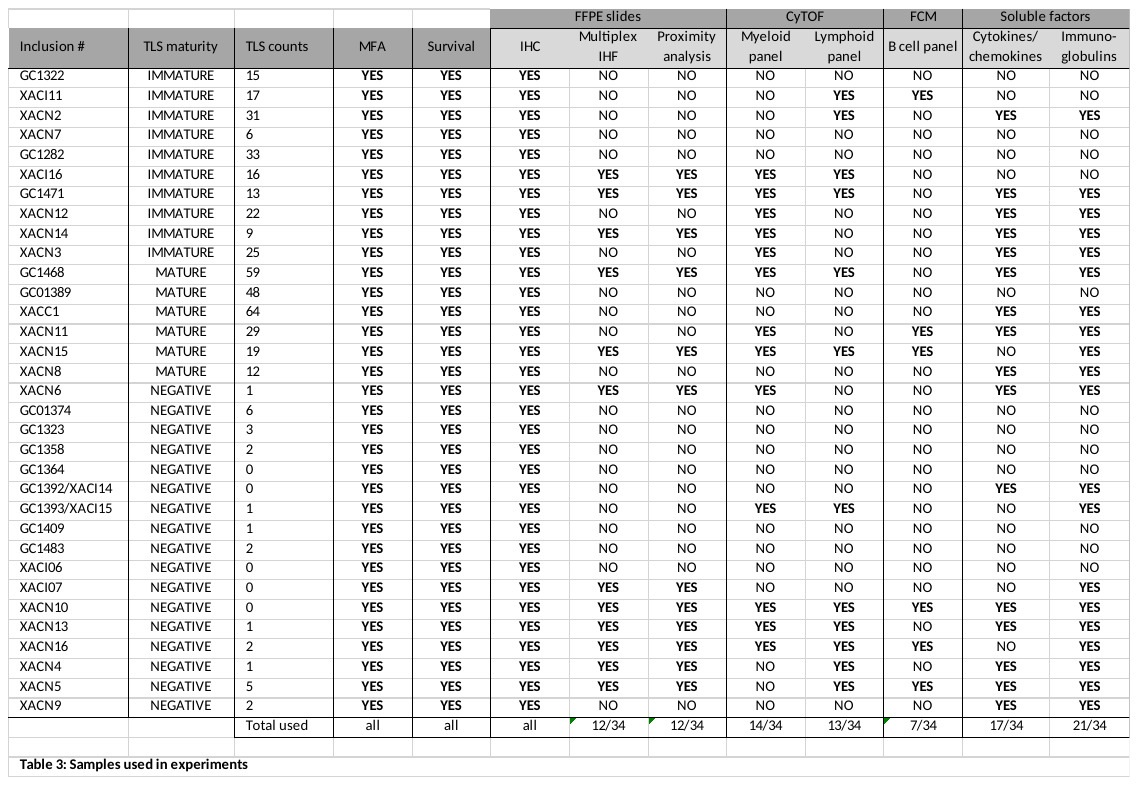
